# Regional Vulnerability of Cardiac Chambers to Radiotherapy: A Multi-Omics Perspective

**DOI:** 10.1101/2025.10.13.681218

**Authors:** Cecilia Facchi, Ardiansah Nugroho, Sami Al-Othman, Hanan AbumanhalMasarweh, Syed Murtuza Baker, Leo Zeef, Gerard M. Walls, Dandan Xing, Izobelle Morrell-Neal, Katharine King, Alexandru Chelu, Sukhpal Prehar, Duncan Forster, Mihaela Ghita-Pettigrew, Marilena Hadjidemetriou, Alicia D’Souza, Luigi Venetucci, Karl T. Butterworth, Elizabeth J. Cartwright, Kaye J. Williams

## Abstract

The heart is highly vulnerable to radiotherapy (RT)-induced injury, leading to molecular and structural remodeling collectively termed radiation-induced cardiac toxicity (RICT). Although several biological pathways have been implicated, the regional, cardiac-specific molecular responses to radiation exposure remain incompletely understood. Here, a multi-omics approach was adopted to longitudinally characterise the unique responses to radiation of the heart base (including ventricular base and right atrium), or the heart apex. Ventricular base irradiation induced a cardiomyopathy phenotype, with pronounced molecular perturbations in metabolism and electrical conduction, while changes related to tissue structure were predominant following apex-directed RT. In the right atrium, irradiation drives fibrotic tissue remodelling, leading to an increased propensity for atrial fibrillation, underpinned by changes in sarcomere organisation. This study represents a comprehensive characterisation of differential spatiotemporal radiation effects in the heart and highlights biological and functional pathways that are potentially clinically actionable for cardiac radioprotection and monitoring.

## INTRODUCTION

Cardiac radiotoxicity encompasses a spectrum of radiation-induced injuries to cardiac structures, commonly observed following thoracic radiotherapy (RT) for malignancies such as breast and lung cancer^1^. Despite technological advancements in RT enabling more precise tumour targeting^2,3^,this can result in radiation-induced cardiac toxicity (RICT), wherein the heart develops various pathologies in the months or years post-treatment. The initial trigger is thought to be endothelial damage that promotes inflammation with subsequent remodelling leading to chronic pathological change^4^. Within weeks, the side effects of radiation doses to the heart manifest as arrhythmias, cardiomyopathy^5^, atherosclerotic cardiovascular disease and valve disease^6^. Higher radiation doses, combined cardiotoxic systemic anticancer agents, and the presence of pre-existing cardiovascular disease can increase the occurrence of RICT^7^. Recent clinical evidence has established a strong association between cardiovascular outcomes and the specific substructures contained within the irradiated volume of the heart^8,9^, underscoring the need for substructure-specific dosimetric assessment and sparing strategies. *In vivo* radiation either at the base of the ventricle or at the apex has been correlated with a reduction in function and histological changes^10,11^. Several preclinical studies have investigated the potential pathophysiological mechanisms underlying RICT, including the importance of the substructures impacted by radiation^12^. However, a comprehensive understanding of the molecular and functional changes over time that determine RICT outcome is lacking^13^. This hinders development of risk models that could be used to develop clinically relevant cardioprotective strategies.

Among potential drivers of RICT, the cardiac conduction system^14–16^ is the least characterised at the molecular and cellular level. The association between RT and the occurrence of conduction defects has been investigated in different oncological circumstances^17^. Adams et al. demonstrated that up to 75% of long-term Hodgkin lymphoma survivors treated with RT had detectable conduction defects after a median follow-up of 14.3 years^18^. In clinical studies, the base of the heart has been reported to be the most vulnerable to RT, leading to poor survival in patients with lung cancer^19^. This anatomical region includes the aortic root, the sinoatrial node (SAN) and the atrioventricular (AV) node within the right atrium, and the origin of the pulmonary veins on the left atrium. Atkins et al. identified specific associations between various arrhythmia subtypes (e.g. atrial fibrillation (AF), supraventricular arrhythmia and ventricular tachycardia) and cardiac substructure radiation dose in lung cancer patients^20^. However, the timing and molecular radiopathology underpinning these observations remains poorly characterised^21^.

In this study, we adopted a longitudinal small animal experimental approach to identify changes that could predict later dysfunction, adopting both functional and multiomics assays. To mimic RT effects in different cardiac regions, we delivered 16 Gy radiation to either the heart base or the apex, as previously described^10^. Using spatial transcriptomics, we found that radiation led to differentially expressed genes (DEG) at 10 weeks follow-up, according to the anatomy irradiated. This observation was confirmed with bulk-RNA sequencing and proteomics of myocardial tissue at the final endpoint of 20 weeks, highlighting the unique response of the ventricular base, apex and the right atrium. Region-specific molecular changes were linked to different functional outcomes, underling the relevance of identifying the irradiated area in thoracic cancer patients to tailor monitoring following treatment, and generating new potential targets for radioprotective drug development.

## METHODS

### Murine cardiac irradiation model

Female C57BL/6J aged 12 weeks (16-20gr; Envigo, UK) were housed in standard housing conditions for laboratory animals (12-hour light-dark cycle, controlled temperature of 19–22°C and humidity of 40–65%) and provided with a standard diet and water ad libitum. Animals were randomly assigned to treatment groups. All experimental procedures were carried out in accordance with the Home Office Guidance on the Operation of the Animals (Scientific Procedures) Act 1986.

Animals were anaesthetised with 2% isoflurane and were irradiated with X-rays (220 kV,13mA) at a dose rate of 2.67 Gy/min on a small-animal radiation research platform (SARRP; Xstrahl Life Sciences, UK). Briefly, cone beam CT (60kV, 0.7mA) collected 360 projections over 1 minute and it was used to reconstruct the 3D image of the whole mouse to identify the heart. Muriplan software was adopted to contour the region of interest, either the base or the apex of the heart to deliver the radiation. A total dose of 16Gy was delivered to the specific substructure of the heart using a 3 × 9 mm collimator, as previously described^10^. Mice weights were monitored throughout the experiment and remained within tolerated weight loss of < 15%.

### *In vivo* analysis-echocardiography and unconscious electrocardiogram

*In vivo* analysis and monitoring of cardiac function was performed 10- and 20 weeks following radiation. Longitudinal transthoracic echocardiography was performed on anaesthetised mice using the VisualSonics Vevo3100® ultrasound system (Fujifilm VisualSonics Inc.). Anaesthesia was adapted to maintain> 450 bpm using <2% isoflurane during recording. Parasternal long-axis videos were recorded over several cardiac cycles to determine ejection fraction (EF) and global longitudinal strain (GLS). Parameter analysis was conducted using Vevo LAB v5.5.1 software. EF was calculated using the LV trace function of the software on the left ventricle long-axis images, while GLS was calculated using the Vevo Strain package (Fujifilm VisualSonics Inc.) after the endocardium was semi-automatically traced and at least three consecutive cardiac cycles were selected^68^. Observers were blinded to the experimental groups during data acquisition.

Analysis of mouse cardiac electrophysiology was performed by adopting an unconscious electrocardiogram (ECG) 1,5,10 and 20 weeks after radiation. Mice were anaesthetised (2% isoflurane) and placed in a supine position on a platform. Surface three-lead ECG was recorded with subcutaneous 2-gauge electrodes attached to the two front paws and the back right paw. The recording was performed with PowerLab/4SP with an ML136 Dual Bio amplifier (ADInstruments). ECG parameters and intervals were identified using Lab Chart 8 software. QTc interval was corrected using Framingham equation^69^.

### Murine electrophysiological analysis

Mice were heparinised with an intraperitoneal injection of Heparin 100 units and euthanized after 10 minutes using cervical dislocation. Excised hearts were cannulated via the ascending aorta with a 22G Langendorff perfusion cannula. Hearts were hung vertically and perfused with oxygenated Tyrode’s solution at 37°C. ECG electrodes were placed in contact with the left atrium and the LV apex, while a pacing electrode was placed on the right atrial appendage. The Mapping Lab EMS64-USB-1003 was used for pseudo-ECG recording and pacing protocols were delivered using the Mapping Lab VCS-3001 Stimulator. The function of the sinus node was evaluated by measuring the sinus node recovery time (SNRT), which was assessed by pacing the atrium for 100 beats at a cycle length of 120ms. The first spontaneous P wave was identified and used to determine the SNRT, which was corrected by subtracting the sinus cycle value. PR interval was calculated during SNRT pacing. Anterograde AV nodal function was assessed using an S1-S1 protocol, pacing the atria at progressively decreasing cycle lengths. The minimal cycle length required to maintain 1:1 AV conduction identified the Wenckebach paced cycle length (WBCL). Programmed right atrial stimulation was performed at a cycle length of 120 ms to determine the AV effective refractory period (AVERP).

Right atrial S1-S11 burst pacing was performed to induce atrial arrhythmias. After recording at baseline, Carbachol (CCH-Merk) was perfused in the tissue and after 10 minutes the burst pacing protocol was repeated. A dose of 1µM was adopted to promote AF phenotype as previously described^70^.

### Bulk-RNA sequencing

RNA-sequencing was performed on murine atrial and ventricular tissue from sham control, apex-irradiated and base-irradiated heart. Twenty weeks after receiving radiation, mice were euthanized and hearts were quickly extracted. Cardiac tissues were dissected, separating the right atrium from the whole ventricle. Base and apex of the ventricle were isolated from both ventricles, considering the 3mm collimator measurement to collect the direct-irradiated tissue. Dissected tissue was immediately snap-frozen and preserved at −80°C. For base and apex irradiated tissue, sham-irradiated tissue was collected from 4 mice while irradiated tissue was collected from 5 mice. Right atrium tissue was dissected from 3 sham-irradiated mice and 4 irradiated hearts. Ventricle and atria RNA was extracted using RNeasy Mini kit and RNeasy Fibrous Tissue Mini kit (Qiagen), respectively, according to manufacturer guidelines. Total RNA (typically 0.025-1µg) was submitted to the Genomic Technologies Core Facility at the University of Manchester. Expanded methods for RNA processing and sequencing are available in the Supplemental Methods section. The output data was used by the bioinformatics facility at the University of Manchester to perform the analysis. Unmapped paired-end sequences from an Illumina HiSeq4000 / NovaSeq 6000 sequencer were tested by FastQC (http://www.bioinformatics.babraham.ac.uk/projects/fastqc/). Sequence adapters were removed, and reads were quality trimmed using Trimmomatic_0.39^71^. The reads were mapped against the reference mouse genome (mm10/GRCm38) and counts per gene were calculated using annotation from GENCODE M25 (http://www.gencodegenes.org/) using STAR_2.7.7a^72^. Normalisation, Principal Components Analysis, and differential expression were calculated with DESeq2_1.40.2^73^. Adjusted p-values were corrected for multiple testing (Benjamini and Hochberg method). Heatmaps were drawn with complexHeatmap v2.12.1^74^. Gene ontology enrichment was studied using clusterProfiler v4.8.3^75^, using p adjusted of <0.1 as cut-off. Ingenuity pathway analysis (IPA, Qiagen) was adopted to perform the networks analysis using as cut off p adjusted (padj) of <0.1 and fold change between 1 and −1.

### Spatial transcriptomics

Ten weeks post radiation, one control mouse, one irradiated at the base and one at the apex were killed by a regulated procedure. Hearts were immediately extracted, washed in Phosphate buffered saline (PBS), fixed in 4% paraformaldehyde and processed to be embedded in wax. Samples were oriented according to the coronal axis and 5µm sections were obtained using Paraffin Microtome RM2255 (Leica) to obtain a 4-chamber view section. Samples were prepared using Visium Spatial Gene Expression for formalin-fixed & paraffin-embedded FFPE kit (protocol CG00040710x Genomics) according to the manufacturer’s guidance. Complete description of sample preparation and sequencing are reported in the Supplemental Methods section. For the downstream analysis, we adopted the Squidpy package in Python. Firstly, we started with quality control (QC) on spots and genes as previously reported for single cell genomics data^76^. We filtered out spots with read counts of less than 300 reads and genes that were expressed in less than 10 spots. We then merged these QC passed spots into a single anndata object for all our further downstream analysis. We first log-normalized (library size normalization) the merged dataset and then computed the neighbours and the UMAP. For clustering, we used the Leiden clustering algorithm and project it on the UMAP for visualization^77^. We applied Wilcoxon test to identify marker genes for identified clusters to then annotate the spots. We then calculate spatially variable genes, genes that have high spatial correlation patterns of expression and vary along the spatial distribution of the tissue structure using Moran’s I, which is the correlation in a signal’s intensity among nearby locations in space^78^. Visualisation and DEG analysis were performed in cellxGene software (CZ CellxGene Discover), using VIP tool. Cluster 6 and 7 have been clustered as erythroid-like cells and adipocytes respectively; therefore, changes in the abundances could be correlated with different presence of residues of blood and pericardial fat in the specimen and therefore they were excluded from the DEG analysis. Cluster 8 has less than 50 cells in total among the samples so it was excluded from the analysis.

### Mass spectrometry analysis

Samples were dissected as described for RNA sequencing analysis and preserved at −80°C. Three biological replicates were performed for each treatment across each cardiac region. Detailed protocols for protein digestion, peptide preparation, and mass spectrometry analysis are available in the Supplemental Methods section.

### Statistical analysis

Statistical analysis and plot data were performed using Prism v9.0 (GraphPad Software, Inc.). Data were initially tested for normality (Shapiro-Wilk). Data are presented as mean ± SEM according to normality analysis, as stated in figure legends. Outlier analysis was performed before the application of statistical comparison. Unless otherwise stated, a p < 0.05 (95% confidence) was considered statistically significant. One-way ANOVA with post hoc multicomparison analysis (Tukey’s) or non-parametric test (Kruskal–Wallis test followed by Dunn’s multiple comparisons) was used to compare significance among 3 or more groups. Comparison between two groups was performed using a T-test for normally distributed data.

### Data availability

Open-access datasets are available from ArrayExpress (www.ebi.ac.uk/arrayexpress) with accession numbers E-MTAB-14630 (for bulk RNA sequencing) and E-MTAB-14646 (for Visium Spatial transcriptomics).

## RESULTS

### Spatial resolution of the effect of radiation on cardiac cell populations

We implemented spatial transcriptomics (ST) to define the molecular profile of different anatomical cardiac regions at 10 weeks post-irradiation. Validation of the approach was performed using markers reported in the cardiac spatial transcriptomic literature to identify cardiac cell subtypes (Supplementary Figure 1)^22^. We identified 1837, 1703, and 1773 spots in apex-irradiated (n=1), base-irradiated (n=1) and sham-irradiated (n=1) tissues, respectively, suggesting homogenous sequencing coverage of the samples (Figure 1a-b). After cluster identification, we categorised each cell population, distinguishing the cell type and correlated biological pathways, using EnrichR software.

**Figure 1:**
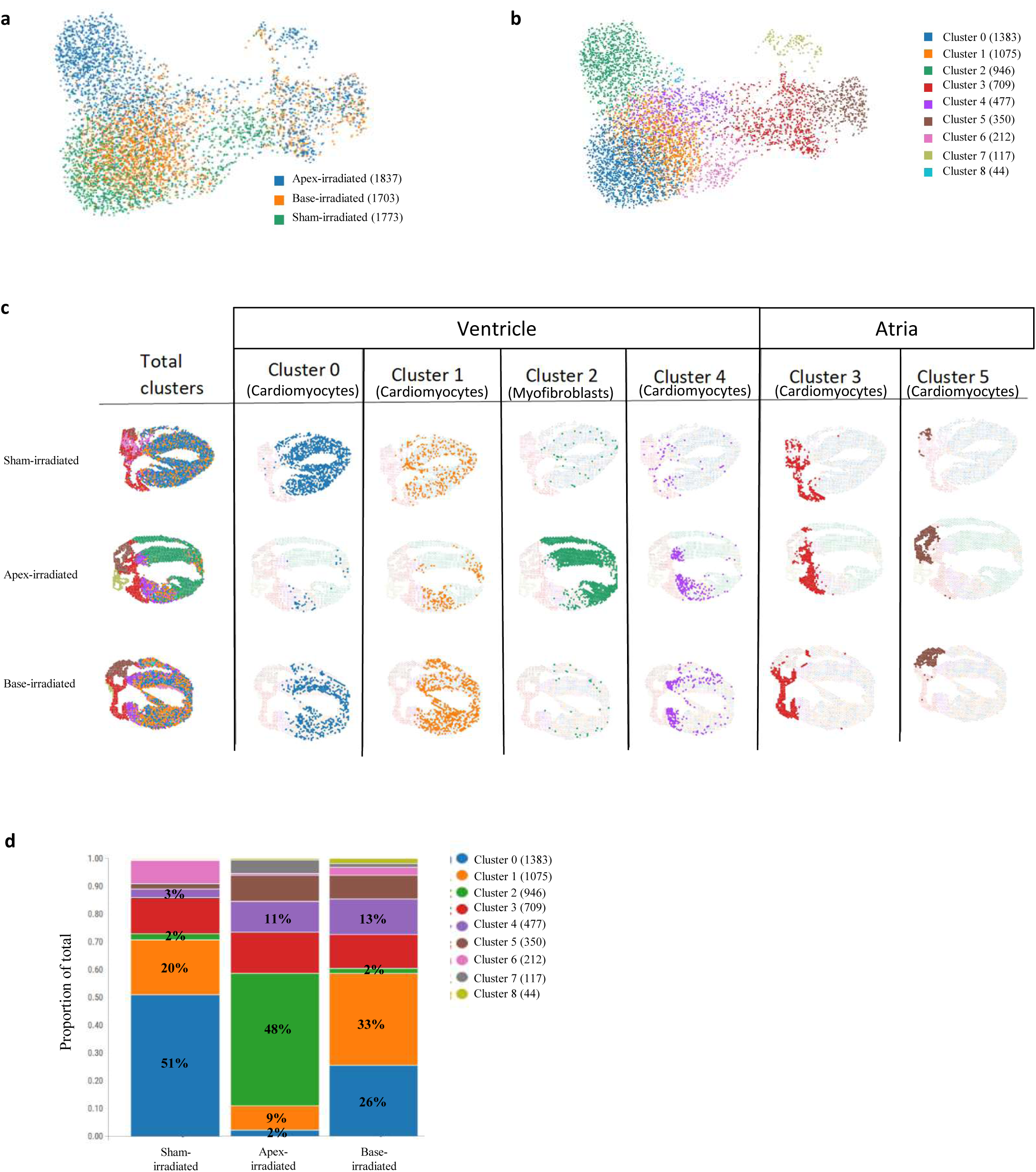
Spatial resolution of radiation effect on cardiac cell populations. **a-b**, Uniform manifold approximation and projection (UMAP) embedding of 5313 spots delineate the distribution according to treatment (**a**) and to the gene expression profile of the different clusters (**b**). In brackets, the number of dots according to samples and clusters classification. **c**, Spatial mapping of cell-type populations in sham-irradiated (n=1 animal), base-irradiated (n=1 animal) and apex-irradiated (n=1 animal) samples. Cluster 0,1,2 and 4 were anatomically identified within the ventricular part of the heart, while clusters 3 and 5 were identified in the atria. **d**, Abundance plot with relative proportion of cell populations according to cell clusters in each sample.

We identified distinctive transcriptional profiles for atrial and ventricular cardiomyocytes^23^, confirming the unique nature of these two populations and highlighting the substantial heterogeneity in cardiomyocyte populations^24^. We identified 3 subpopulations of ventricle cardiomyocytes, represented by clusters 0,1, and 4 (Figure 1c) with no differences in these cluster distributions between the left and right ventricles. These clusters showed sarcomere encoding gene enrichment (Table 1). Cluster 0 represented the conventional ventricular cardiomyocyte population, indicated by the expression of canonical genes connected with metabolic pathways for cardiac muscle contraction^25^, such as fatty acid degradation and respiratory electron transport (e.g. *Hadha*; *Eno3*; *Ckm*) (Table 1). Clusters 3 and 5 represented atria cardiomyocytes and both clusters showed enrichment of muscle contraction-related pathways. However, Cluster 5 presents markers highly associated with right atrium identity, such as *Smad6 and Bmp10* and gene enrichment pathway analysis highlighted upregulation of adrenergic signallingcorrelated genes (*Ryr2; Tpm2; Tnnt2; Creb3l2; Gnas; Atp2a2; Atp1a1; Atp1b1; Slc8a1*)^26^, suggesting a specialisation of those cells in contraction and electrical impulse propagation. Further spatial visualisation of the remaining clusters (clusters 6,7 and 8) can be found in Supplementary Figure 2.

**Table 1:** Gene-based classification of cluster’s cell types and biological pathways. Description of the 9 identified clusters, with relative cell population identity (“cell type identity” column) using the first 10 marker genes for each population (“gene signature” column). PanglaoDB Augmented database was used for this analysis. Pathway analysis was performed with the first 100 markers genes per cluster using KEGG Human 2021 and Reactome 2022 database. The first hits are reported for each cluster. Both analysis were performed using EnrichR as platform. For a complete table of cluster markers, please see Supplementary material.

| Cluster number | Cell type identity (PanglaoDB Augmented) | Main biological pathways (KEGG, Reactome, Bioplanet) |
| --- | --- | --- |
| 0 | Cardiomyocytes | 1. Metabolism<br>2. Respiratory electron transport, ATP biosynthesis<br>3. Tricarboxylic acid (TCA) cycle and respiratory electron transport |
| 1 | Cardiomyocytes | 1. Dilated cardiomyopathy<br>2. Adrenergic signaling in cardiomyocytes<br>3. Citrate cycle (TCA cycle) |
| 2 | Myofibroblasts | 1. Vitamin C in brain<br>2. Bone mineralization regulation<br>3. Angiotensin-converting enzyme 2 regulation of heart function |
| 3 | Cardiomyocytes | 1. Fas signalling in cardiomyocytes<br>2. Corticosteroids and cardioprotection<br>3. YAP- and TAZ-stimulated gene expression |
| 4 | Cardiomyocytes | 1. Hypertrophic cardiomyopathy<br>2. Cardiomyocyte hypertrophy<br>3. Sarcomere in cardiomyocytes in dilated cardiomyopathy |
| 5 | Cardiomyocytes | 1. Muscle contraction<br>2. Regulation of complement cascade<br>3. Complement cascade |
| 6 | Erythroid-like and erythroid precursor cells | 1. Haptoglobin binding<br>2. Haemoglobin beta and alpha binding<br>3. Oxygen carrier activity and binding |
| 7 | Adipocytes | 1. Fatty acid metabolism<br>2. Metabolism<br>3. triglyceride biosynthesis |
| 8 | Smooth muscle cells | 1. Apelin signaling pathway<br>2. Vascular smooth muscle contraction<br>3. TCA cycle |

The spatial distribution of clusters highlighted the unique response of cardiac tissue to radiation according to the irradiation site (Figure 1c). In the sham-irradiated sample, ventricular cardiomyocyte cluster 0 represented 51% of the overall spots (Figure 1d). Upon irradiation of the apex, the dominant ventricular population became cluster 2, identified as myofibroblasts, distributed throughout the ventricle. Interestingly, this population was almost absent in the base-irradiated and shamirradiated heart. Pathway enrichment analysis confirmed characteristic fibroblastrelated processes, such as organisation of the extracellular matrix (*Col15a1; Mmp2; Fn1; Col3a1; Col4a2; Adam15; Col4a1*), as they are responsible for scar/fibrosis maturation^27^. The distribution pattern of cluster 2 highlights changes manifested beyond the irradiated volume. This observation is extended with base-irradiation causing shifts within ventricular populations whereby the proportion of cluster 0 to cluster 1 shifts from 51 vs 20% in sham to 26 vs 33% in base-irradiated samples. Genes in cluster 1 correlate to diabetic cardiomyopathy development in the ventricle (e.g.*CD36, Tnni3*)^28,29^, suggesting that cardiomyocytes assume distinct pathological phenotypes dependent on the site of irradiation.

A specific radiation-associated subpopulation of cardiomyocytes was identified in the ventricular base, represented by cluster 4, which increased from 3% up to 11-13% in irradiated tissues. The main representative genes were *Myl3, Myl7* and *Myl4*, which correlate with pathological rearrangement of the sarcomere^30^. Interestingly, both apex- and base-irradiated tissues showed an enlargement of this population, despite the location of the radiation, again suggesting that phenotypic change extends beyond the directly irradiated tissues.

Together these analyses highlight considerable and distinct changes in the distribution of cell populations, according to the irradiated site, and those changes manifest beyond the directly irradiated tissue.

### Spatial transcriptomics revealed unique molecular changes across the ventricle according to the radiation area

Irradiation of either the heart base or apex caused differential gene expression across the ventricle with changes observed beyond irradiated volume. Focusing on clusters 0,1,2,4, we performed DEG analysis to compare ventricular gene expression in sham versus base and apex irradiated samples. Differential responses to radiation were highlighted by the specific DEG in apex-irradiated (n=29, Supplementary Material S1) and base-irradiated hearts (n=19, Supplementary Material S2) (Figure 2a-b). DEG following apex-irradiation included *CD74, Fabp4* and *Slc25a4*, whilst *Mb, Mybpc3* and *Tcap* were dysregulated in the ventricle following base-irradiation (Figure 2c). To identify specific dysregulated pathways, we adopted an enrichment pathway analysis using Kyoto Encyclopedia of Genes and Genomes (KEGG). Apexirradiation led to an enrichment of pathways correlated with the immune response (Figure 2d top). Particularly, myocarditis was observed, confirming what has been previously reported as a generic RICT phenotype^31^. Interestingly, ventricle tissue following heart base-irradiation showed a striking cardiomyopathy-related enrichment, followed by cardiac contraction and metabolism (Figure 2d bottom). To correlate the genes to specific pathways, we performed gene-enriched k-mean cluster analysis in STRING. Biological pathways (BP) of the Gene Ontology (GO) database highlighted changes in pathways correlated to antigen presentation, collagen extracellular matrix and hypertrophy in ventricular tissue following apexirradiation (Figure 2e), while processes connected to sarcomere organisation and oxidative phosphorylation were altered in the base-irradiated sample (Figure 2f).

**Figure 2:**
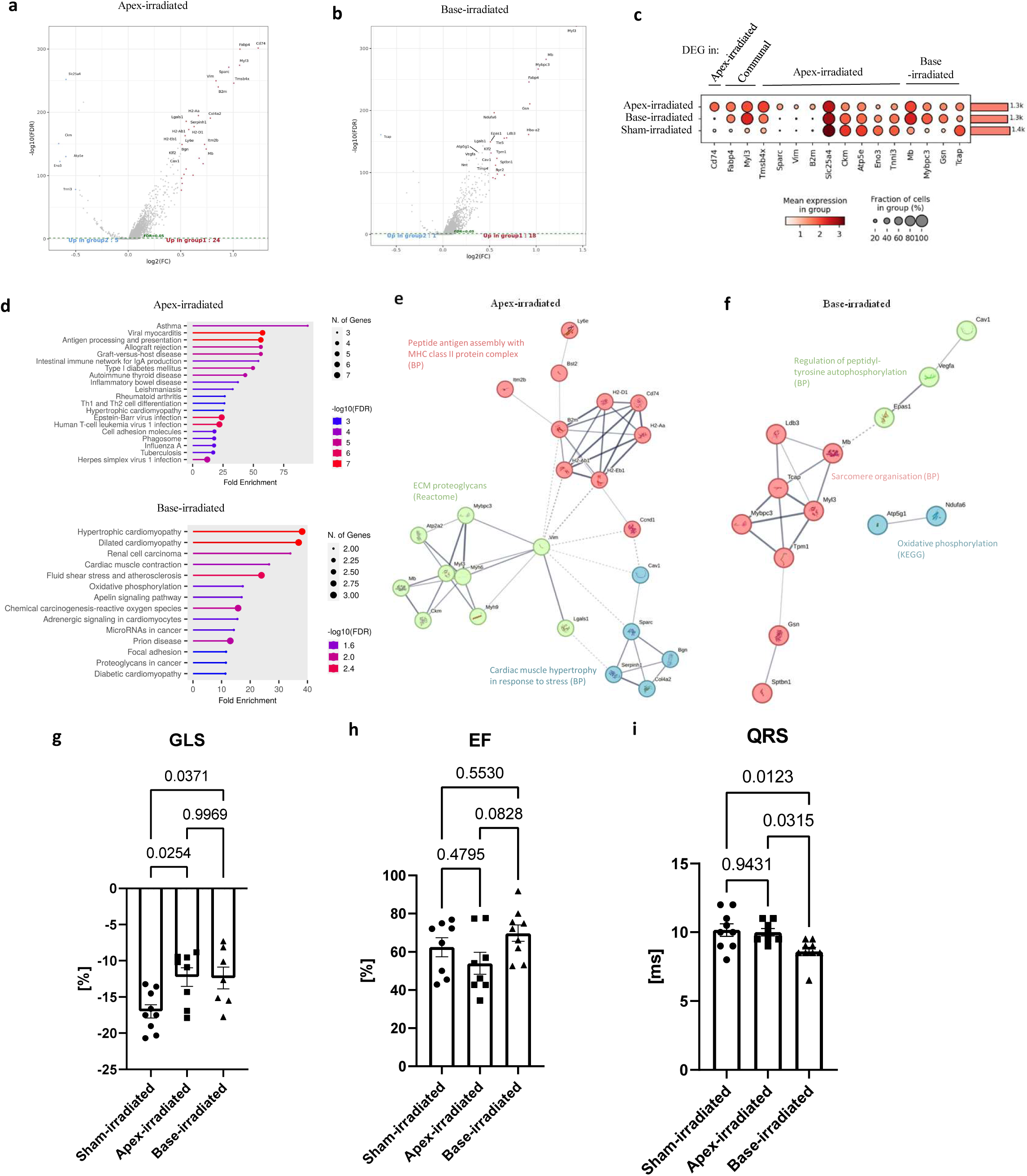
At ventricular level, ST revealed unique molecular changes according to radiation area, leading to dysfunction and electrical conduction alteration. **a-b**, DEG analysis (absolute log2(FC)>0.5, FDR<0.05) represented by volcano plot for apex- (a) and base- (b) irradiated tissue. Welch’s t-test identified 24 up- and 1 down-regulated genes in apex-irradiated heart compared to sham-irradiated. In base-irradiated tissue, 18 up- and 1-downregulated genes were identified. Upregulated genes are identified in red and downregulated genes in light blue. **c**, Dot plot with relative expression of the dysregulated genes in both apex and base irradiated heart. A value of 200 −log10(FDR) was adopted as cut off. The colour of dots indicates level of expression of each marker, while the size displays the percentage of cells expressing it. **d**, Lollipop charts of the first 15 pathways of the dysregulated genes were identified using ShinyGo 0.80 based on KEGG database for apex-irradiated (top) and base-irradiated tissue (bottom). FDR cut off= 0.05 was used for this analysis. **e-f,** STRING plot showing functional interrelationship of dysregulated genes in the apex- (n=29; **e**) and base-irradiated (n=19; **f**) tissue. K-Mean analysis was performed using 3 clusters/sample. Identification of functional patterns was performed with Biological Pathway (BP; Gene Ontology). Kyoto Encyclopedia of Genes and Genomes (KEGG) or Reactome databases were adopted for the classification in case of missing BP identification, as reported in the figure. The analysis was performed with STRING 12.0. Disconnected dots were hidden from the network. Inter-cluster edges are represented by dashed-lines. **g,** Echocardiographic analysis of cardiac performance using GLS 10 weeks after radiation. Sham-irradiated: n=9 biological replicates; apex-irradiated= 8 biological replicates; base-irradiated= 7 biological replicates. **h**, EF quantification from echocardiography analysis. Sham-irradiated: n=8 biological replicates; apex-irradiated= 8 biological replicates; base-irradiated= 9 biological replicates. **i**, Quantification of QRS interval via unconscious ECG recording. Sham-irradiated: n=9 biological replicates; apex-irradiated= 8 biological replicates; base-irradiated= 8 biological replicates. For all graphs, data are represented as mean ±SEM. One-way AVONA was adopted for normally distributed data. P values are reported in the graphs and p<0.05 was considered statistically significant. MHC: Major histocompatibility complex; ECM: Extracellular matrix; GLS=Global Longitudinal Strain; EF=Ejection fraction; ECG: electrocardiogram

Considering the differential molecular changes after radiation at the ventricle level, we investigated the impact of site-specific tissue irradiation on ventricular systolic function and ventricular electrical conduction *in vivo* at the same time point (10 weeks after radiation). Echocardiography identified severe impairment of cardiac performance associated with incorrect myocardial deformation demonstrated by the reduction in global longitudinal strain (GLS) in both groups of radiation-treated mice (base-irradiated and apex-irradiated) (Figure 2g). Ejection fraction (EF) remained unaffected at this timepoint (Figure 2h). Longitudinal ECG analysis showed that mice exposed to 16 Gy irradiation exhibited alterations in ventricular depolarization, as shown by the shortening of QRS (Figure 2i). Interestingly, these changes were observed only in the base irradiated group, suggesting spatial specificity of the radiation effects on the cardiac conduction system. No changes were observed in other ECG intervals (Supplementary Table 1).

As we observed that pathways correlated with sarcomere organisation were affected in the base-irradiated group, we investigated if this was related to changes in the expression of ion channels and molecules involved in electrical conduction which could result in the observed ECG changes. Limited alterations in the expression of K^+^ channels were observed; however, variations in the abundance of genes coding for Na^+^/Ca^2+^ handling ion channels, such as *Scn5a*, *Cacna1c, Slc8a1, Ryr2* and *Atp2a2* were identified (Supplementary Figure 3). Increased expression in *Ryr2*, which is a crucial regulator of Ca^2+^ sarcoplasmic reticulum release^32^, was observed in DEG analysis in parallel with a significant decrease in *Tcap* expression, suggesting a prominent disruption in the propagation of the conduction potential, considering that *Tcap* is required for the maintenance of T-tubule structure and function^33^.

### Bulk-RNA sequencing and proteomics at 20 weeks after radiation highlighted local changes in irradiated tissue

We next assessed the longer-term impact of radiation on the specific regions of the heart at the functional level and we found that the decreases in GLS identified at 10 weeks post-irradiation were preserved at 20 weeks after radiation (Supplementary Figure 4). At this later time point, we found that there was also a reduction in EF in the apex-irradiated mice. Therefore, we investigated the possible exacerbation of pathological pathways at the transcriptional level. Region-specific bulk-RNA sequencing analysis of irradiated tissue confirmed that the different locations of the ventricle exhibited differential remodelling 20 weeks after radiation. In apex-irradiated tissue, statistically upregulated genes were *CD163* and *Mpg*, both known to have a proatherogenic role^34,35^ (Figure 3a). Irradiation at the base of the ventricle led to upregulation of genes correlated with vascular remodelling^36^, such as *Angpt2*, and higher expression of MHCII genes, including *H2-Eb1, H2-Aa, H2-Ab1*^37^ (Figure 3b). BP of GO analysis of apex-irradiated ventricle dysregulated genes (n=1092 with padj<0.1, Supplementary Material S3) showed enriched collagen-organisation pathways (Figure 3c), with an upregulation in cellular proliferation, while extracellular signal-regulated kinase (ERK)1 and ERK2 pathways were the main dysregulated biological pathways in base-irradiated tissue (n=156 with padj<0.1, Supplementary Material S4) (Figure 3d). Ingenuity Pathway Analysis (IPA)-based analysis highlighted the main genes correlated with the relevant pathways, suggesting new potential candidates for therapeutic targets (Supplementary Figure 5). In particular, collagen and TNF-related cellular movement genes were observed dysregulated in apex-irradiated tissue, while base-irradiated ventricle showed pathways correlated with dysregulation of ERK and G protein-coupled receptor (GPCR), which are essential for cardiac function, including heart rate and contraction. Moreover, associations between Angpt1/Angpt2 pathway and collagen formation and focal adhesion kinase were identified among IPA networks in base-irradiated mice. These analyses confirmed that the base and apex of the ventricle have a unique transcriptional remodelling profile after radiation, leading to area-specific molecular changes that could potentially underpin the differential clinical toxicities related to specific cardiac structures.

**Figure 3:**
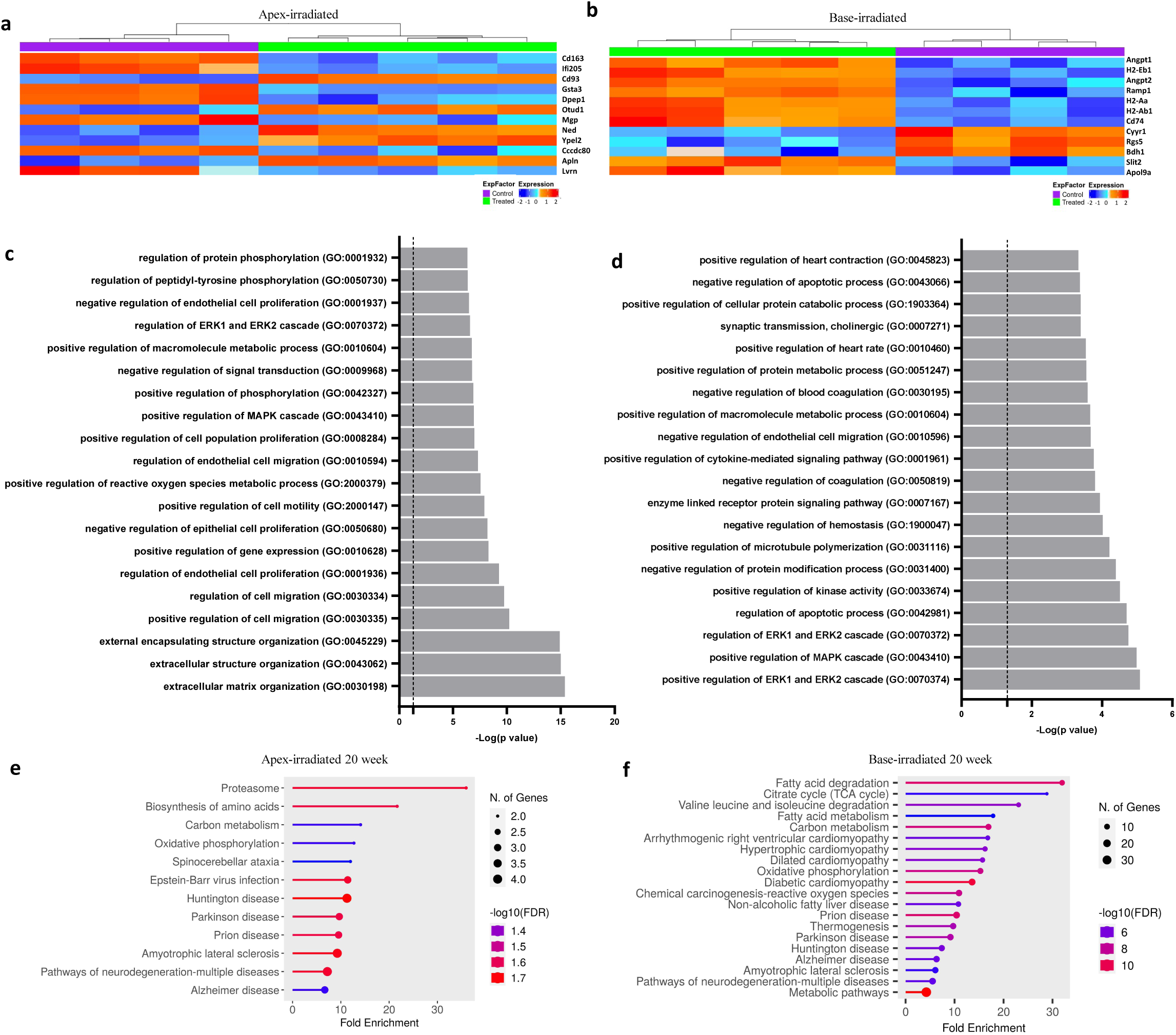
Bulk-RNA sequencing and proteomics analysis 20 weeks after RT highlighted local changes in irradiated tissue. **a-b**, RNA-sequencing-derived heat maps of the first 12 most differentially expressed genes listed by pAdjusted values between sham-irradiated and apex (**a**) or base-irradiated (**b**) tissues. **c-d**, Bar plot of BP GO analysis of 20 most significant biological pathways from RNA-sequencing analysis of dissected tissue of apex (**c**) and base-irradiated (**d**) tissues. Dotted lines represent FDR < 0.05 of p value< 0.05. **e-f**, Lollipop plot of dysregulated proteins (Apex-irradiated tissue: 41; Base-irradiated tissue: 444) highlighting KEGG-based biological pathways. FDR cut off= 0.05 was used for this analysis.

To identify potential protein markers correlated with the specific changes at the tissue level, proteomics analysis was performed at 20 weeks after radiation from each dissected irradiated cardiac region. Apex-irradiated tissue proteomics highlighted 41 differentially expressed proteins (Supplementary Material S5), while we identified pronounced remodelling in base-irradiated tissue compared to shamirradiated tissue (Supplementary Material S6), as demonstrated by the 444 differentially abundant proteins (Supplementary Figure 6a-b). Venny-based analysis of differentially expressed proteins highlighted unique changes between base- and apex-irradiated tissues (Supplementary Figure 6c). KEGG-based pathway analysis identified proteasome and metabolism changes as the main dysregulated pathways in apex-irradiated hearts (Figure 3e). Analysis of the changes in the base-irradiated tissue identified that several of the dysregulated proteins correlated to metabolic pathways, such as fatty acid degradation and citrate cycle, suggesting remodelling at the metabolic level of the cardiomyocytes, aligning with the RNA-Sequencing data (Figure 3f). To identify protein-protein interactions in base-irradiated tissue, we adopted STRING K-Mean analysis of the 100 top-upregulated proteins, which highlighted molecules specifically involved in muscle structure development and sarcomerogenesis (*Actn2, Myl9, Ttn*), suggesting that cellular structure was still remodelling at this stage (Supplementary Figure 7).

### Radiation-induced gene changes in the right atrium led to perturbation of conduction properties and development of fibrosis

In our study, regions of right atrial tissue were also irradiated when targeting the heart base. The right atrium is extensively involved in the initiation and propagation of cardiac depolarisation to the remainder of the heart. To investigate the role of radiation on arrhythmia development, we investigated the molecular changes in cardiomyocytes of the right atrium 10 weeks following irradiation of the heart-base. Firstly, we demonstrated that cardiomyocytes were the major cell type in the right atrium, indicated by the presence of unique atrial cardiomyocyte markers, including *Nppa* and *Myl7* (Supplementary Figure 8). Atria-specific analysis of 38 DEG (Supplementary Material S7) indicated higher expression of pro-fibrotic genes (Figure 4a-b), such as *Ccn2*, in the base-irradiated sample, suggesting a role for those cardiomyocytes in the formation of fibrosis in the atrium. KEGG-based pathway enrichment analysis identified pathways correlated to cardiomyopathy development (Figure 4c) and STRING K-mean analysis of dysregulated proteins highlighted changes in myofilament organisation, contributing to the development of a dilated cardiomyopathy molecular phenotype (Figure 4d). Next, we investigated the transcriptome in the dissected right atrium using bulk-RNA sequencing. BP analysis of right atrium dysregulated genes (n=513 with padj<0.1, Supplementary Material S8) showed that the main dysregulated pathways were linked to extracellular matrix organisation and collagen fibres, suggesting that the right atria tissue could be transitioning to a pro-fibrotic state at this time point (Figure 4e). Specific dysregulation of genes associated with collagen formation and electrical conduction properties was observed by IPA (Supplementary Figure 9a-b). Moreover, Upstream Regulator Analysis (URA) in IPA identified abnormal morphology and enlargement of the heart chamber as final overall effects of pathways dysregulated after radiation, supporting the hypothesis of pathological atrial remodelling (Supplementary Figure 9c). Proteomics-based analysis of dysregulated genes (Supplementary Material S9) highlighted that purine metabolism was the main dysregulated pathway in the irradiated tissue (Figure 4f), supporting a change in tissue organisation as alteration of this pathway has been correlated with development of dilated cardiomyopathy^38^.

**Figure 4:**
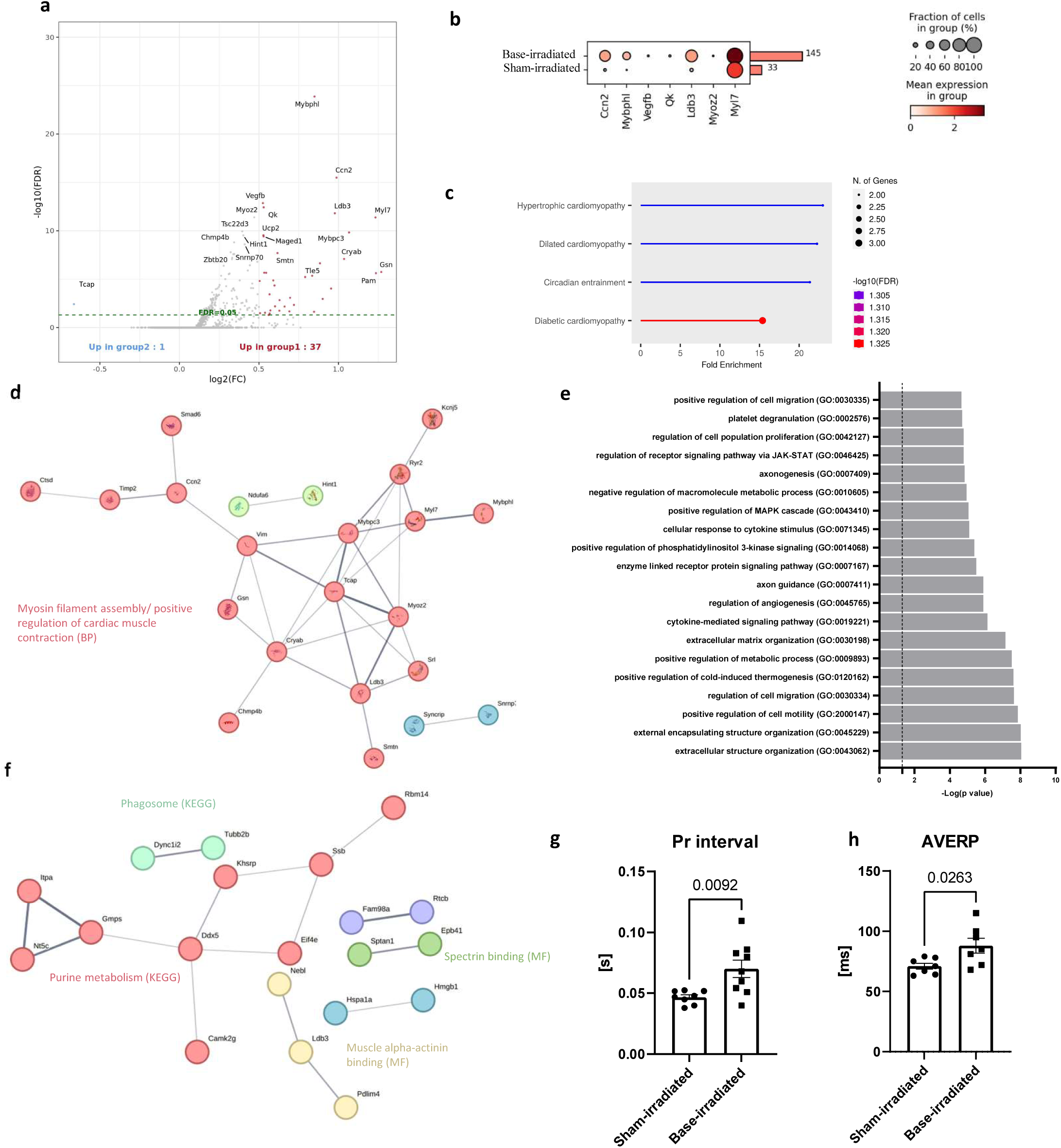
Right atrium remodelling could explain the impact of RT on the conduction system. **a**, ST–derived volcano plot of DEG analysis (absolute log2(FC)>0.5, FDR<0.05) using Welch’s t-test, which identified 38 dysregulated genes in right atrium tissue 10 weeks after RT at the base of the heart. **b**, Dot plot visual representation of statistically significant up- and down-regulated DEG in ST. −log10 (pvalue) cut-off was adopted to screen the statistically significant abundant proteins to represent in the graph. **c,** KEGG-based lollipop plot of significant biological pathways of 38 DEG of ST. **d,** STRING plot with K-mean analysis of dysregulated DEG in ST. Identification of functional patterns was performed with BP. No pathway was correlated to green and blue clusters due to the limited number of belonging proteins. **e**, Bar plot of first 20 BP GO pathways of bulk-RNA sequencing of right atrium 20 weeks after RT. FDR < 0.05 of p < 0.05. **f**, Proteomic-based STRING plot after K-mean analysis of 53 statistically dysregulated proteins. Among them, 33 proteins resulted as disconnected nodes. Identification of functional patterns was performed with Molecular function (MF; Gene Ontology). KEGG was adopted for the classification in case of missing MF identification. Purple and dark blue clusters were not correlated to any biological pathway. **g,** Pr interval quantification during ex vivo pacing. Unpaired t-test was adopted with normally distributed data. Sham-irradiated: n=6 biological replicates; base-irradiated: n=9 biological replicates. **h,** AVERP data were normally distributed and unpaired t-test was adopted. Sham-irradiated: n=7 biological replicates; base-irradiated: n=7 biological replicates. For all graphs, data are represented as mean±SEM. P value is reported in graph.

To further investigate the effect of radiation-mediated tissue remodelling of the right atrium, we performed pacing to investigate SAN and atrioventricular (AV) node function. Surprisingly, no statistical difference was observed in SAN activity (Supplementary Figure 10a), between base and sham-irradiated hearts, suggesting an absence of modification of the function of this structure. However, the autonomic signal-independent PR interval was significantly longer in the base-irradiated group compared to sham-irradiated mice (Figure 4g), suggesting a slowing of AV node conduction between the atria and ventricles. Remarkably, the Atrioventricular Effective Refractory Period (AVERP) was prolonged in base-irradiated animals (Figure 4h), suggesting AV node dysfunction, which was supported by potential changes in Wenckebach cycle length (WBCL) (Supplementary figure 10b). Altogether, these data suggested that heart base radiation involving atrial tissue can cause alteration in the function of the conduction substructures within the right atrium, such as AV node, leading to an impairment of impulse propagation from the AV node.

### Radiation at the heart base leads to higher susceptibility to AF

To specifically explore the effect of heart-base irradiation on the development of atrial arrhythmias, we investigated the presence of an atrial arrhythmogenic phenotype. Langendorff-based *ex vivo* pacing highlighted an increased atrial tachycardia phenotype in the base-irradiated group (Figure 5a), with 22% (2/9) of the mice developing AF (Figure 5b). After the addition of the acetylcholine receptor agonist Carbachol (CCH), 89% of the base-irradiated mice exhibited AF during the pacing protocol. In contrast, only 2/8 sham-irradiated mice showed AF episodes after CCH infusion. Among the base-irradiated animals, we observed the development of sustained AF, both at baseline (Figure 5c) and after CCH perfusion (Figure 5d), suggesting an intrinsic change in the structure of the tissue that could create a substrate for arrhythmia^39^.

**Figure 5:**
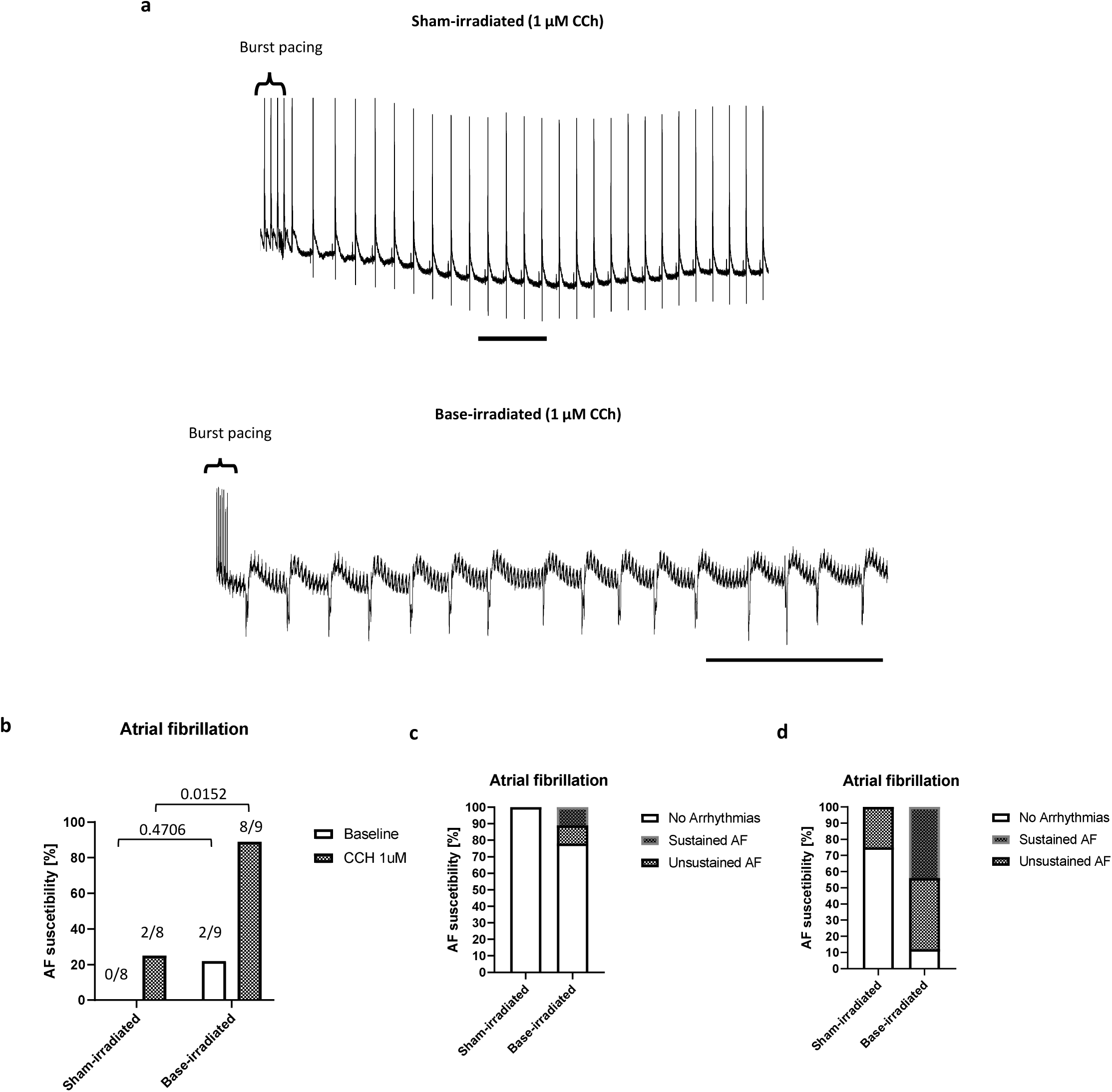
Impact of RT at the base of the heart on AF occurrence. **a**, Representative ex-vivo pacing traces after burst pacing for AF induction in sham-irradiated (top) and base-irradiated heart (bottom), after perfusion with 1µM of CCH. Scale bars= 1 Sec. **b**, AF episodes quantification at baseline and with 1µM CCH perfusion. Fisher’s exact test was applied. Exact numbers of AF episodes are reported in the graph in relation to the total number of analysed mice. **c-d,** Classification of the nature of AF in bar plot at baseline (**c**) and after CCH perfusion (**d**). b-d Sham-irradiated: n=8 biological replicates; base-irradiated: n=9 biological replicates. AF=atrial fibrillation; CCH: Carbachol.

## DISCUSSION

Despite an increased understanding of the pathological basis of RICT, optimal guidelines for follow-up and the prevention of cardiac side effects after radiation remain to be fully described^40,41^. This is due to a lack of understanding of how the variety of pathologies linked to RICT, such as myocardial fibrosis, conduction abnormalities and coronary diseases, manifest and the classical view that the heart is uniformly radiosensitive^42^. Emerging evidence has clearly demonstrated that regions of the heart have differences in radiosensitivity. These concepts are further supported by our findings as we have demonstrated functional, electrophysiological and transcriptional differences following radiation treatment of cardiac-specific subvolumes (Figure 6). These data could potentially explain the differences observed in the clinical landscape, where radiation at the heart base has been correlated with worse survival in lung cancer patients^43,44^.

**Figure 6:**
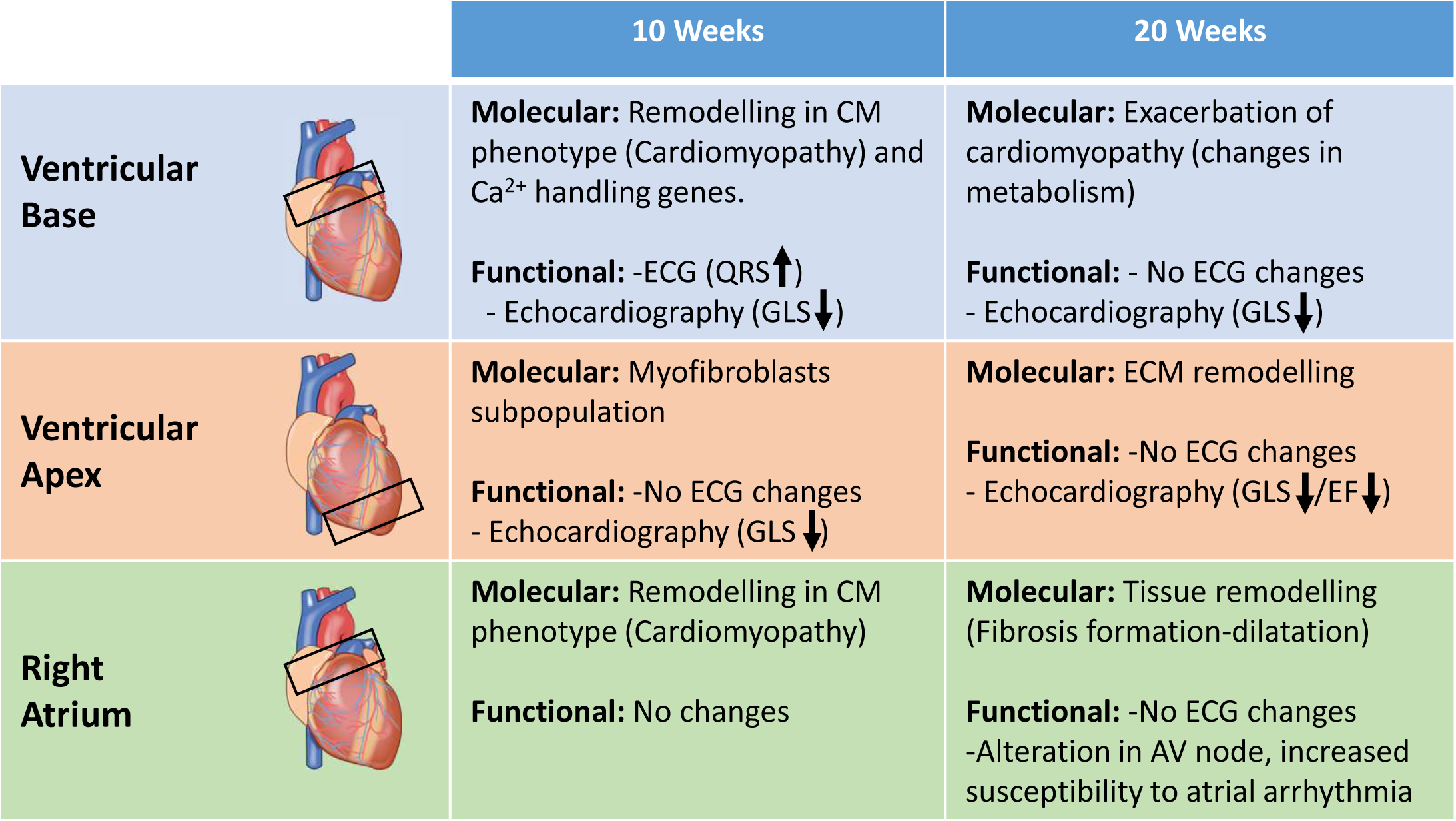
Schematic representation of the main characteristics for each region according to functional, transcriptomics and proteomics analysis.

Base-irradiated tissue showed molecular and cellular changes towards cardiomyopathy 10 weeks after radiation, which could represent an opportunity for toxicity early detection in patients. ST and proteomics highlighted changes in sarcomere protein abundance and oxidative phosphorylation, which has been previously correlated with increased muscle mass due to cardiomyopathy development^45^. Unbalanced fatty acid metabolism and alteration of mitochondria activity have been previously correlated with fibrosis pathogenesis after radiation and cardiomyopathy development^46–48^. Particularly, the switch to glycolysis is strongly correlated to fibrosis formation and limitation of cardiac regeneration. A prevalence higher than 10% has been estimated for radiation-induced cardiomyopathy^49^.

In apex-irradiated tissue, identification of myofibroblasts upon cluster analysis indicates significant, potentially pathogenic, remodelling. In terms of function, myofibroblasts could limit the contractility capacity of the ventricle but support a prolonged function of the organ in time. Dreyfuss previously reported a decrease in cardiac function in apex-irradiated tissue due to fibrosis using a higher dose of radiation (40 Gy)^11^; we identified this phenotype in our mouse model, with a significant reduction of EF in apex-irradiated mice. However, the EF range was still within the physiological values (apex-irradiated EF mean: 70%; Supplementary Figure 4), suggesting a limited effect due to the lower dosage RT protocol adopted in this study or additional compensatory changes. The latter may be linked with myofibroblast-mediated extracellular matrix organization and/or macrophage-like properties exhibited by myofibroblasts^50^. Proteomics analysis suggested an upregulation of proteasome machinery, widely reported in extracellular matrix reorganization orchestrated by myofibroblasts, possibly reflecting an attempt of the tissue to overcome the radiation-induced damage.

RNA sequencing data suggested that apex-irradiated tissue is characterised by extracellular matrix remodelling correlated genes, as demonstrated by the GO analysis. Among the most upregulated genes, we identified *CD163, Gsta3 and Mgp,* which are associated with calcification and subsequent development of atherosclerosis^51–53^. Those markers are predictors of plaque phenotype but also cardiovascular-specific mortality and incident heart failure, so specific blood analysis in thoracic cancer patients where the apex of the heart is irradiated could allow the preventive identification of early pathological signals for vessel diseases.

Previous clinical studies have reported that conduction abnormalities are another side effect of RT. Arrhythmias are reported as appearing within 2 months of the end of the RT^5^. Longitudinal analysis of ECG data demonstrated that according to the specific radiation-targeted regions, the conduction system is differentially affected over time, with a particular effect when the radiation is directed to the base of the ventricle. Shortening of the QRS complex was previously observed by Zhang et al.^14^, where they identified this phenomenon in mice treated with whole-heart radiation after 6 weeks. Here, we showed QRS interval shortening specifically after radiation to the base of the ventricle at 10 weeks, proposing a different sensitivity of the anatomical regions. Anatomically, the heart-base radiation volume includes the passage of the impulse from the AV node to the Bundle of His before its propagation to the bundle branches in the distal ventricle. ST analysis revealed the accumulation of *Ryr2* and disruption of the t-tubule network genes, coupled with the downregulation of *Tcap*, which is a key molecule involved in Ca^2+^-induced Ca^2+^ release^33^, suggesting changes in the depolarization of the ventricle. T-tubules are essential invaginations of the cytoplasm that allow the uniform spread of depolarisation in ventricular cardiomyocytes^54^. These findings suggest that irradiation can strongly modulate Ca^2+^ handling at the ventricular level after base radiation^55^, which could lead to physiological alterations. However, changes in function and conduction were modest and non-progressive despite the significant molecular changes, as shown by the 20-week time point. This suggests that the myocardium has the potential to compensate to some extent the structural remodelling and support normal function, which is in line with clinical data, where 70% of the ECG abnormalities can return to normal 6 months after RT^5^. However, early monitoring of ECG in patients who complete RT could predict late dysfunction as we observed in our animal model.

Sinoatrial (SAN) and atrioventricular (AV) nodes reside in the right atrium and they are frequently co-exposed during radiation to the base of the heart. In our study, we investigated if the direct tissue remodelling of the right atrium could lead to the alteration of cardiac conduction. Cardiomyocytes are still present in this substructure, but they clearly showed dysregulation of genes correlated with the propagation of Ca^2+^ sparks (e.g. *Tcap* and *Ryr2*). Ryr2 upregulation at the protein level has been recently reported 60 weeks after radiation by Feat-Vetel et al. in parallel with a drastic presence of fibrosis in the right atrium, suggesting that our analysis could have identified early molecular defects that lead to an exacerbated phenotype at later timepoint^56^. Moreover, cumulative deposition of collagen could lead to a slow deteriorating phenotype, which precipitates an environment conducive to AF arrhythmogenesis in the right atrium. We identified higher expression of *Ccn2* in the right atrium after radiation^57^. Increased levels of *Ccn2* have been previously reported in cardiac tissue in C57BL/6J mice receiving a single X-ray dose of 16 Gy to the heart 40 weeks after radiation^58^. The early detection of this marker could signify a higher susceptibility of right atrium tissue to radiation and *Ccn2* could contribute to the tissue remodelling, potentially predicting fibrotic development at a later time point and being a potential therapeutic target. This is supported by the findings from the proteomics analysis, where base-irradiated right atrium tissue showed purine metabolism pathway variation, which is essential for ATP production, and activation of this pathway has been correlated with the early phases of myocardial infarction in ischemic myocardium^59^, suggesting a potential involvement of this metabolic rearrangement in the response to radiation. This reorganisation at the tissue level could lead to the dysfunction that we observed in the SAN-AV nodes.

The hyperactivity of Ryr2 has been correlated to increased AF susceptibility in cardiac disease^60^, and we observed significant upregulation of Ryr2 in our baseirradiated model. To support this concept, we adopted carbachol, which specifically activates Ryr2^61^ and increases the utilization of intracellular Ca^2+^ SR stores during parasympathetic dominance^62^. Ex vivo pacing identified increased prevalence of arrhythmias in base-irradiated hearts which was markedly enhanced, and sustained, with carbachol treatment, validating the increased susceptibility of the atria to arrhythmias after radiation^63^. These data indicate that radiation-induced arrhythmias can arise following irradiation of the heart base, potentially due to impact on the SAN, AV and overall conduction system after RT. Further investigation would be required to understand the relative contribution of right-atria-driven effects to overall propensity for radiation-induced arrhythmia, recognising that the potential contribution of dose to the left pulmonary vein has not been directly addressed in these studies^64,65^.

This work has a direct relevance to patient care. Physiological and functional markers that can distinguish the effect of RT and the relative affected cardiac region are limited in the literature. Altered GLS has been previously reported as a marker of radiation dysfunction in base-irradiated with different cardiac radiation regimes^10,66^. GLS is a good early marker but does not necessarily differentiate between radiation sites. The risk of conduction abnormalities and arrhythmogenic phenotype align most strongly with heart base irradiation, tailoring monitoring towards these pathologies and recognising that there may be enhanced risk of detrimental outcomes for patients with pre-existing conditions. Similarly, for patients where apex ventricular regions lie within the target radiation field, monitoring for vessel damage markers may be the favoured approach. Our highlighting of molecular mechanisms and tissue remodelling has created a multi-omics database as a valuable resource for the future identification of RICT markers and to aid the identification of potential intervention strategies to mitigate detrimental responses and improve patient outcomes. In conclusion, our study provides a deep molecular and functional characterisation of radiation effects on different cardiac sub-regions, in particular, concerning the cardiac conduction system.

This study has limitations, for example, the presented data are derived only from female mice. Small animal analysis and clinical evidence reported that females have a higher incidence of RICT compared to men, especially when they are postmenopausal, highlighting a discrepancy between the sexes^67^. Potential differential responses according to sex should be considered in future evaluations and appropriate follow-up in patients. In this study, we presented a valuable spatial analysis of the effect of RT on different cardiac regions, however, ST resolution is limited as each identified dot corresponds to approximately 20 cells, which could restrict the interpretation of the disposition at a single cardiac cell level. Moreover, the annotation for each dot is the summary of the most abundant population within the dot, losing the information about the remaining cells. Therefore, the possibility of adopting single-cell analysis in parallel to ST would have enabled discrete distinction of cell populations. Moreover, ST was performed on one single plane of each heart, which prevents the comprehensive 3D disposition of the cell populations. Finally, as mentioned, the ex vivo AF pacing protocol did not permit localisation of the origin of AF as the recorded stimulus is produced by both atria.

## ACKNOWLEDGMENTS

This work was supported by Cancer Research UK RadNet Manchester programme [C1994/A28701] (K.W.). The authors would like to thank Dr Stacey Warwood from the Mass Spectrometry facility of the University of Manchester for her valuable help in processing proteomics samples. We also would like to thank the Genomic Technologies Core Facility at the University of Manchester, in particular Mrs Beverley Anderson, for the support with the sequencing process for bulk Rna sequencing and spatial transcriptomics. Finally, the support from the Bioinformatics Core Facility team for Rna sequencing and spatial transcriptomics analysis. Thanks to NIAID Visual & Medical Arts. Human Heart 000228 & Lab Mouse 000279. 07/26/2024 & 9/26/2024. NIAID BIOART Source. bioart.niaid.nih.gov/

## AUTHOR CONTRIBUTIONS

**Cecilia Facchi**: Conceptualization; Data curation; Formal analysis; Investigation; Visualization; Writing - original draft. **Ardiansah Nugroho**: Conceptualization; Investigation. Writing – review & editing. **Sami Al-Othman**: Investigation; Resources. Writing – review & editing. **Hanan Abumanhal-Masarweh**: Investigation; Resources. Writing – review & editing. **Syed Murtuza Baker**: Data curation; Formal analysis; Software. **Leo Zeef**: Data curation; Formal analysis; Software. **Gerard M. Walls**: Conceptualization; Writing - review & editing. **Dandan Xing**: Investigation. **Izobelle Morrell-Neal**: Formal analysis. **Katharine King**: Formal analysis. **Alexandru Chelu**: Software. **Sukhpal Prehar**: Resources **Duncan Forster**: Methodology. **Mihaela Ghita-Pettigrew**: Writing – review & editing. **Marilena Hadjidemetriou**: Resources, Methodology. **Alicia D’Souza**: Resources, Methodology. **Luigi Venetucci**: Supervision. **Karl T. Butterworth**: Writing – review & editing, Methodology, Conceptualization. **Elizabeth J. Cartwright**: Writing – review & editing, Writing – original draft, Supervision, Resources, Methodology, Investigation, Data curation, Conceptualization. **Kaye J. Williams**: Writing – review & editing, Writing – original draft, Supervision, Resources, Project administration, Methodology, Investigation, Funding acquisition, Formal analysis, Data curation, Conceptualization.

## COMPETING INTERESTS STATEMENT

The authors declare no competing interests.

## ABBREVIATIONS

RT: radiotherapy
ECG: electrocardiogram
HF: heart failure
AF: atrial fibrillation
DEG: differentially expressed genes
ST: spatial transcriptomics

## Notes

### Competing Interest Statement

The authors have declared no competing interest.

