## Supplementary Methods, Figures and Table for "Regional Vulnerability of Cardiac Chambers to Radiotherapy: A Multi-Omics Perspective"

### Supplementary Information

### Methods

##### Bulk-RNA sequencing

After the extraction, the quality and integrity of the RNA samples were assessed using a 4200 TapeStation (Agilent Technologies) and then libraries were generated using the Illumina® Stranded mRNA Prep. Ligation kit (Illumina, Inc.) according to the manufacturer's protocol. Briefly, total RNA was used as input material from which polyadenylated mRNA was purified using poly-T, oligo-attached, magnetic beads. Next, the mRNA was fragmented under elevated temperature and then reverse transcribed into first strand cDNA using random hexamer primers and in the presence of Actinomycin D (thus improving strand specificity whilst mitigating spurious DNA-dependent synthesis). Following removal of the template RNA, second strand cDNA was then synthesized to yield blunt-ended, double-stranded cDNA fragments. Strand specificity was maintained by the incorporation of deoxyuridine triphosphate (dUTP) in place of dTTP to quench the second strand during subsequent amplification. Following a single adenine (A) base addition, adapters with a corresponding, complementary thymine (T) overhang were ligated to the cDNA fragments. Pre-index anchors were then ligated to the ends of the doublestranded cDNA fragments to prepare them for dual indexing. A subsequent PCR amplification step was then used to add the index adapter sequences to create the final cDNA library. The adapter indices enabled the multiplexing of the libraries, which were pooled before cluster generation using a cBot instrument. The loaded flow-cell was then paired-end sequenced (76 + 76 cycles, plus indices) on an Illumina HiSeq4000 instrument. Finally, the output data was demultiplexed and BCLto-Fastq conversion was performed using Illumina's bcl2fastq software, version 2.20.0.422.

##### Spatial transcriptomics

Briefly, sections from each heart were collected on the Visium Spatial Gene Expression Slide (PN-2000233). Deparaffinization, Haematoxylin & Eosin (H&E) staining, imaging, and decrosslinking of FFPE tissues were then performed (protocol CG000409-10x Genomics). FFPE stained slides were imaged using Olympus BX63 single slide scanner microscope (Olympus Lifescience). Images were collected on an Olympus BX63 upright microscope using a 20x objective, captured and white balanced using a DP80 camera (Olympus) through CellSens Dimension v1.16 (Olympus). Images were then processed and adopted for Loupe alignment as described below. Then, using CG000407 protocol from 10x Genomics, Visium Mouse Transcriptome Probe Set v1.0 was used for hybridization, ligation and extension steps. After sample cycle determination using qPCR, the library was constructed and sequenced. After the sequencing using Illumina Nova Seq6000, bcl2fastq (Illumina) was used to generate the fastq files, which were then mapped to the respective genome using Space Ranger v.1.2.0 (10x Genomics). The Loupe alignment file, which stores the manual alignment for the fiducial markers (10x Genomics), and details of tissue sample slides were input as prerequisites in Space Ranger.

**Heart tissue preparation and mass spectrometry analysis**

After tissue collection, homogenization was conducted in pre-made RIPA buffer (Sigma-Aldrich) supplemented with protease inhibitor mini tablets (Pierce) in AFA tubes (Covaris) using the LE220-plus focused-ultrasonicator (Covaris). To obtain the protein lysate, samples were centrifuged at 14,000 RCF for 10 minutes, and the supernatant was collected. Proteins were then alkylated and reduced with dithiothreitol (DTT, 5 mM) and iodoacetamide (IAM, 15 mM), respectively. Protein concentration was quantified using a BCA assay (Pierce). Subsequently, 10 µg of total protein was processed using the S-Trap method. Briefly, protein lysates (total volume 50 µL) were acidified with phosphoric acid (12%) and mixed with six volumes of binding buffer (100 mM TEAB in 90% methanol, pH 7.1). Proteins were loaded into S-Trap micro columns (ProtiFi, LLC, USA) by centrifugation (4,000 x g, 2 min, 4°C) and washed four times with the binding buffer. Proteins were then digested with trypsin (0.1 µg/mL) for 1 hour at 47°C. Peptides were eluted from the S-Trap columns using digestion buffer (50 mM TEAB), 0.1% formic acid in water, and 30% acetonitrile in water containing 0.1% formic acid. Peptide samples were desalted using OligoTM R3 beads (Thermo Scientific), stored in 50% acetonitrile in water with 0.1% formic acid in 30% acetonitrile, and dried in a Speedvac (Thermo Scientific).

Samples were run using Orbitrap Exploris™ 480 mass spectrometer. The separation was performed on a Thermo RSLC system consisting of a NCP3200RS nano pump, WPS3000TPS autosampler, TCC3000RS column oven and analytical column (Waters nanoEase M/Z Peptide CSH C18 Column, 130Å, 1.7 µm, 75 µm X 250 mm) configured with buffer A as 0.1% formic acid in water and buffer B as 0.1% formic acid in acetonitrile. A multistage gradient was used varying the percentage of buffer B, as described: 1% to 6% B over 2 minutes, 6% to 18% over 44 minutes, 18% to 29% over 7 minutes and 29% to 65% over 1 minute before washing for 4 minutes at 65%. Then, the multistage gradient was dropped down to 2% B in 1 minute. The analytical column was connected to a Thermo Exploris 480 mass spectrometry system via a Thermo nanospray Flex Ion. Fragmentation data was obtained from signals with a charge state of +2 or +3 and an intensity over 5,000 and they were dynamically excluded from further analysis for a period of 15 sec after a single acquisition within a 10ppm window. Data were processed using Proteome Discover v3.1 (Thermo Fisher).

### Figures

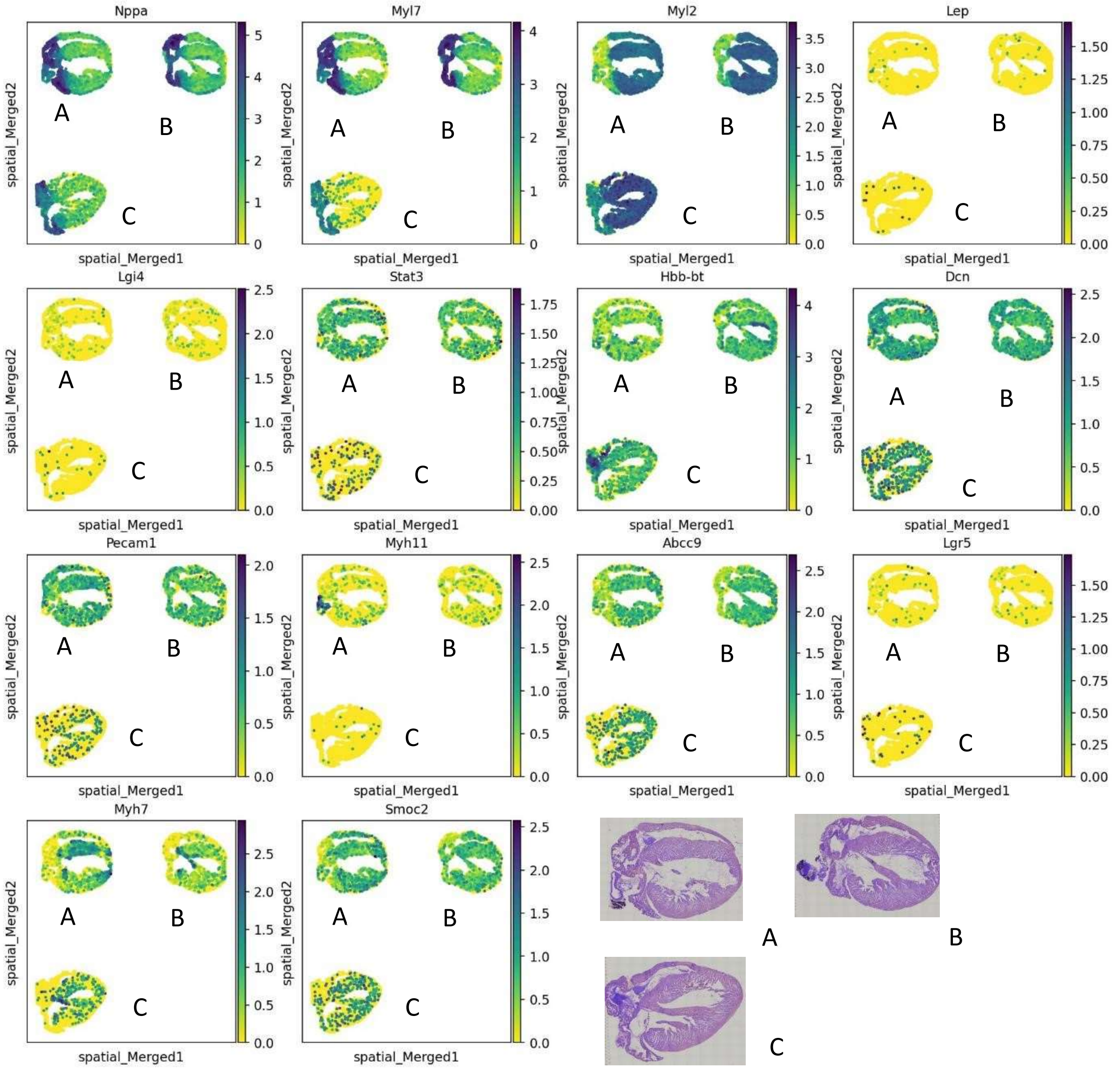

**Supplementary Figure 1. Representative cardiac markers for cell population in ST.** Markers were plotted for each sample and embedding plot was adopted to identify any change in the distribution of the populations. NPPA/myl7: atria cardiomyocytes, Myl2/ MyH7: ventricle cardiomyocytes, Lep: mediastinal fat, Lgi4: neuronal cell, Stat3: macrophages, Hbb-bt: erythrocytes, Dcn: fibroblasts, Pecam1: endothelial cells, Myh11: vascular smooth muscle cells, Abcc9: pericytes, LgR5/Smoc2: cardiac conduction cells. At the bottom of the panel, the corresponding histological specimen stained with Haematoxylin staining. A: apexirradiated; B: base-irradiated; C: sham-irradiated.

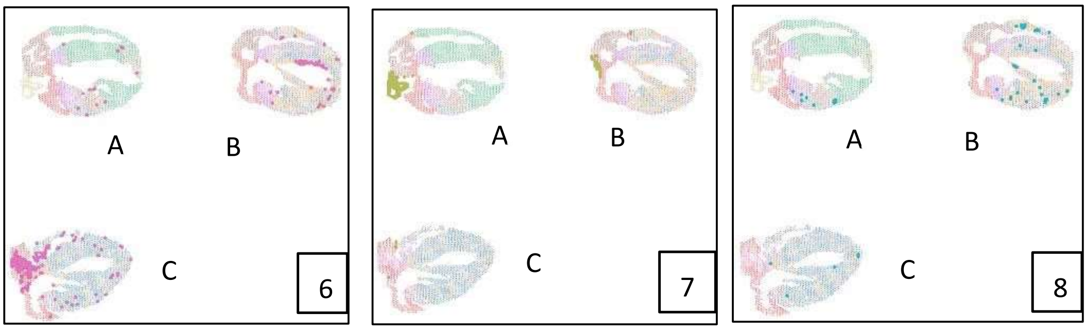


**Supplementary Figure 2. ST clusters 6-7-8 visual distribution**. Additional clusters identified during the cluster analysis. Cluster number is represented in the box. As presented in the main text, Cluster 6 and 7 represented residues of blood and pericardial fat, respectively. Cluster 8 was classified as smooth muscle cells. A: apex-irradiated; B: base-irradiated; C: sham-irradiated.

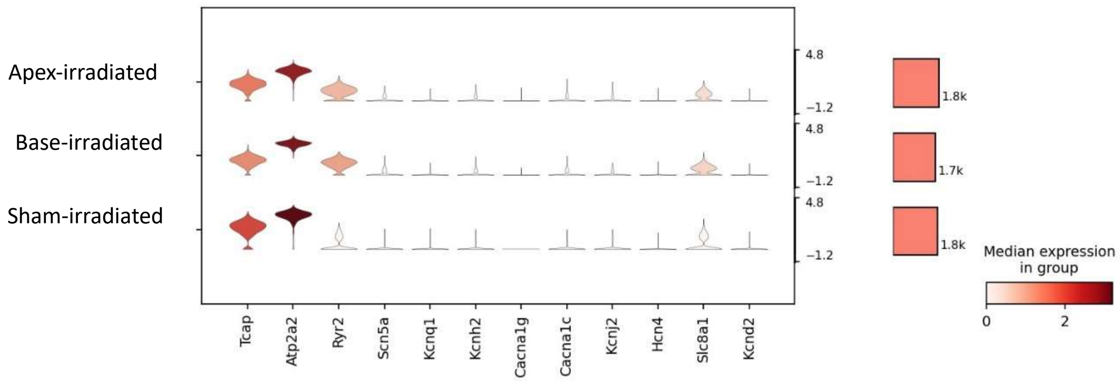


**Supplementary Figure 3. Analysis of conduction-related $Na^{+}$-$K^{+}$-$Ca^{2+}$ ion channels**. Violin plot analysis of ion channels involved in the propagation of electrical impulse in ST at ventricle level. Red bar on the right reported the number of analysed cells/sample in the ventricular clusters. Expression level is represented by the bar on the right.

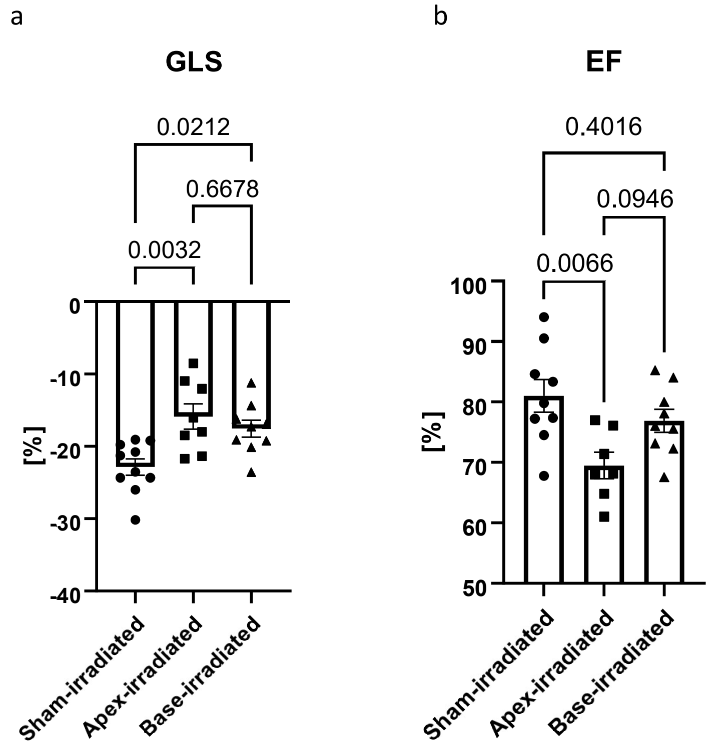


**Supplementary Figure 4. GLS (a) and EF (b) echocardiography parameters 20 weeks after RT.** Data are represented as mean ±SEM. One-way AVONA was adopted for normally distributed data and p value are reported for each comparison. GLS: Sham-irradiated: n=10; apex-irradiated= 8; base-irradiated= 9. EF: Sham-irradiated: n=9; apex-irradiated= 7; base-irradiated= 9.

a

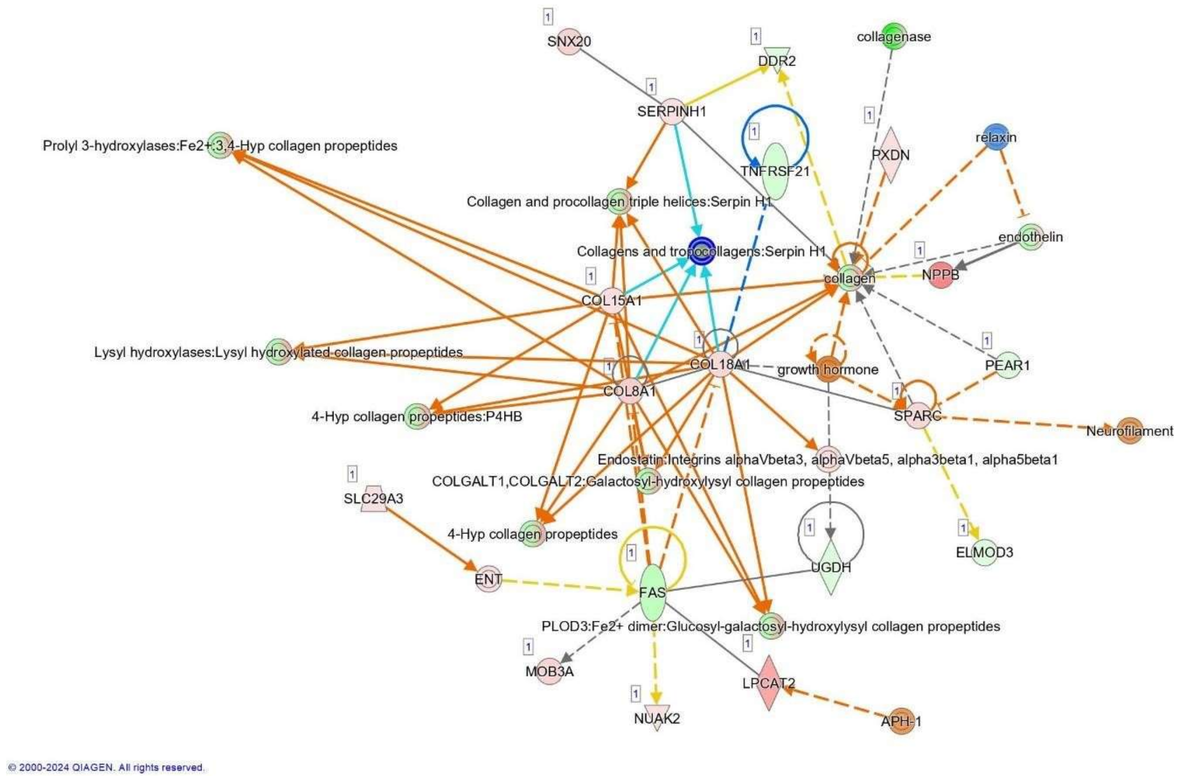
SNX20
DDR2
collagenase
SERPINH1
Prolyl 3-hydroxylases:Fe2+:3,4-Hyp collagen propeptides
PXDN
relaxin
TNFRSF21
Collagen and procollagen triple helices:Serpin H1
endothelin
Collagens and tropocollagens:Serpin H1
NPPB
collagen
COL15A1
Lysyl hydroxylases:Lysyl hydroxylated collagen propeptides
COL18A1
growth hormone
PEAR1
COL8A1
4-Hyp collagen propeptides:P4HB
SPARC
Neurofilament
Endostatin:Integrins alphaVbeta3, alphaVbeta5, alpha3beta1, alpha5beta1
COLGALT1,COLGALT2:Galactosyl-hydroxylysyl collagen propeptides
SLC29A3
4-Hyp collagen propeptides
ELMOD3
UGDH
ENT
FAS
PLOD3:Fe2+ dimer:Glucosyl-galactosyl-hydroxylysyl collagen propeptides
MOB3A
LPCAT2
APH-1
NUAK2
© 2000-2024 QIAGEN. All rights reserved.

b

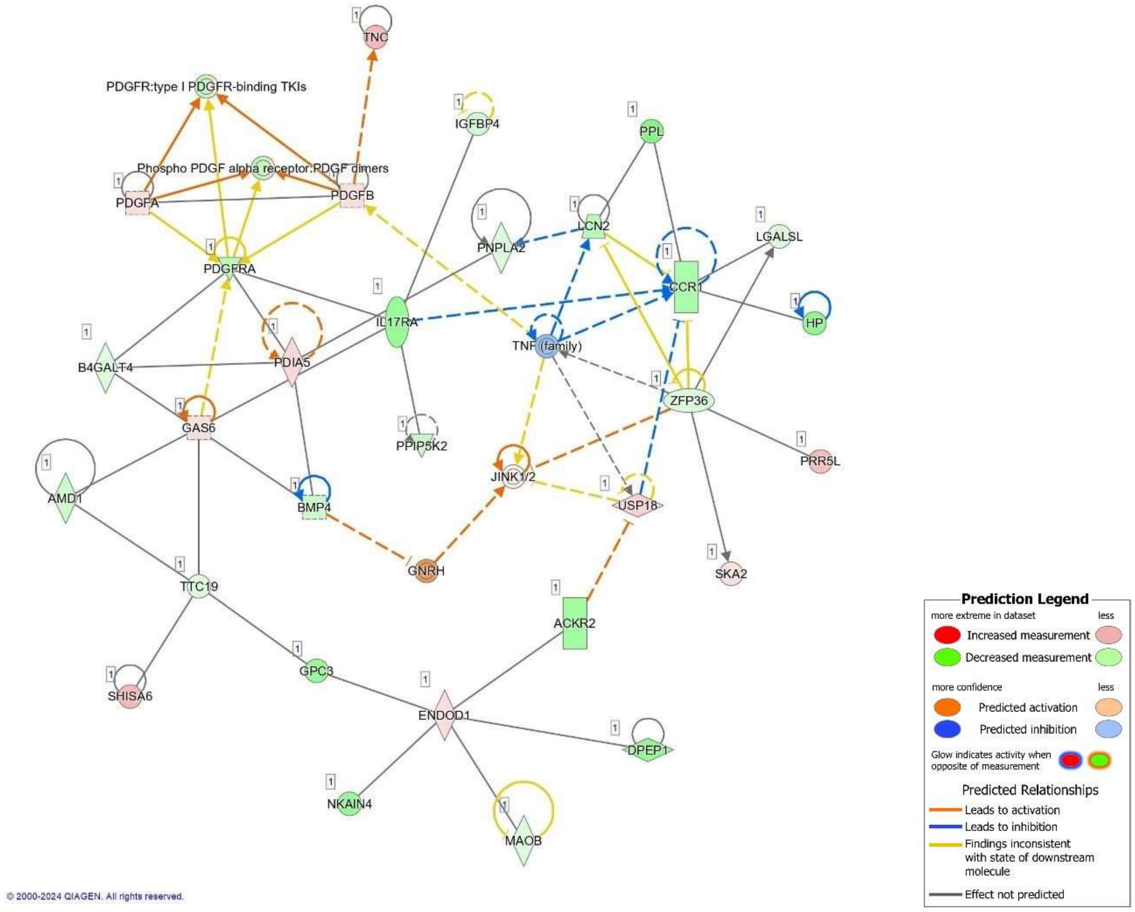
TNC
PDGFR:type I PDGFR-binding TKIs
IGFBP4
PPL
Phospho PDGF alpha receptor:PDGF dimers
PDGFA
PDGFB
LCN2
LGALSL
PNPLA2
PDGFRA
CCR1
IL17RA
HP
TNF (family)
B4GALT4
PDIA5
ZFP36
GAS6
PPIP5K2
PRR5L
JINK1/2
AMD1
BMP4
USP18
GNRH
SKA2
TTC19
ACKR2
GPC3
SHISA6
ENDOD1
DPEP1
NKAIN4
MAOB
Prediction Legend
more extreme in dataset
less
Increased measurement
Decreased measurement
more confidence
less
Predicted activation
Predicted inhibition
Glow indicates activity when opposite of measurement
Predicted Relationships
Leads to activation
Leads to inhibition
Findings inconsistent with state of downstream molecule
Effect not predicted
© 2000-2024 QIAGEN. All rights reserved.

c

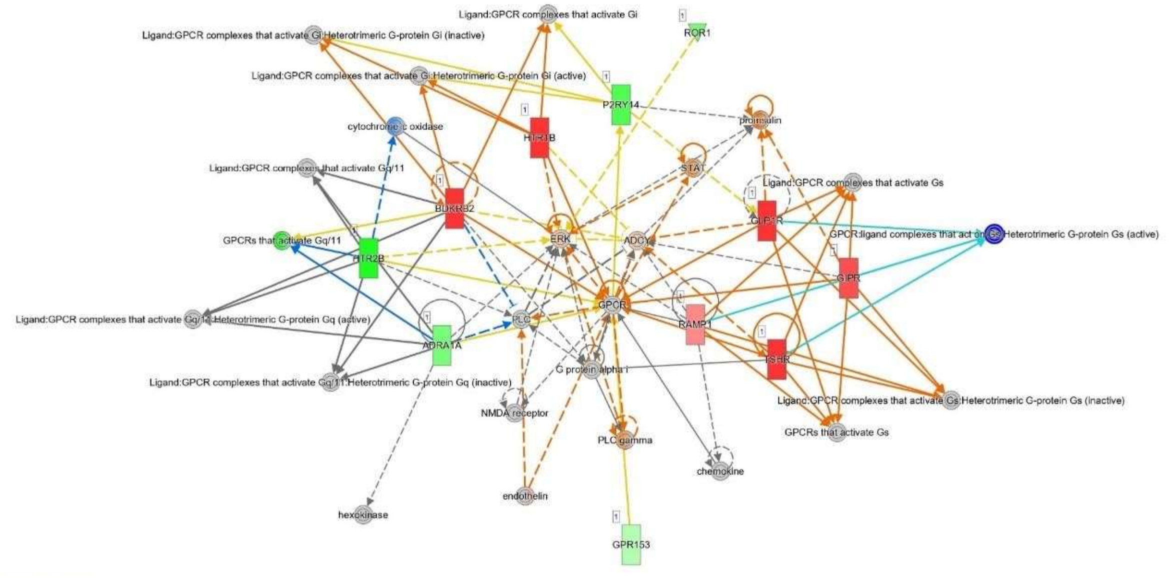


© 2000-2024 QIAGEN. All rights reserved.

d

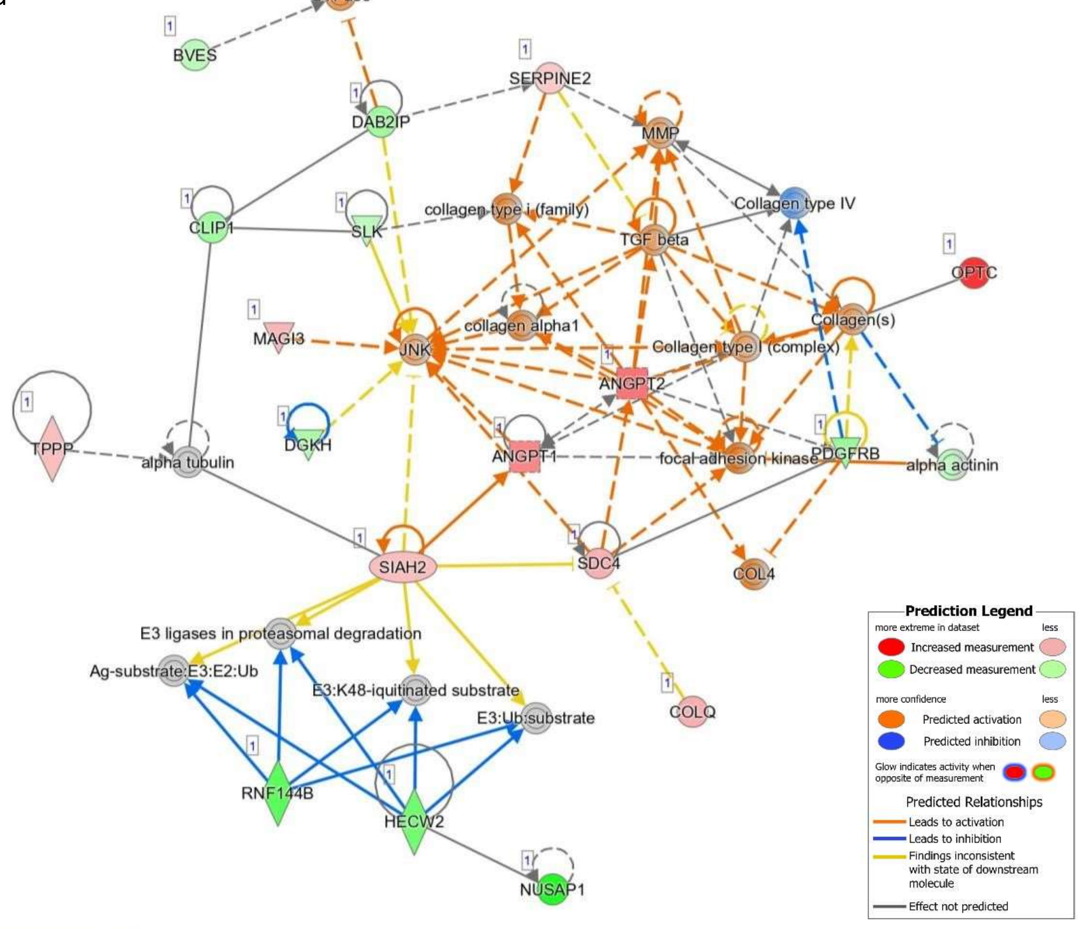


© 2000-2024 QIAGEN. All rights reserved.

**Supplementary Figure 5. RNA-Sequencing analysis of ventricular base and apex using IPA.** Ingenuity Pathway Analysis (IPA) network of genes from RNA-Sequencing filtered using pAdjusted <0.1 dysregulated between sham-irradiated and apex-irradiated (a,b) or sham-irradiated versus base-irradiated tissue (c,d). Network, algorithmically generated based on their functional and biological connectivity, was graphically represented as nodes (genes) and edges (the biological relationship between genes). Expression of genes is reported according to the legend. Solid and dashed lines between genes represent known direct and indirect gene interactions, respectively. The shapes of the nodes reflect the functional class of each gene product: transcriptional regulator (horizontal ellipse), transmembrane receptor (vertical ellipse), enzyme (vertical rhombus), cytokine/growth factor (square), kinase (inverted triangle) and complex/group/other (circle). IPA nomenclature: a: collagen-related genes; b: cellular movements; c: ERK-related signaling; d: Angpt1-2 related processes.

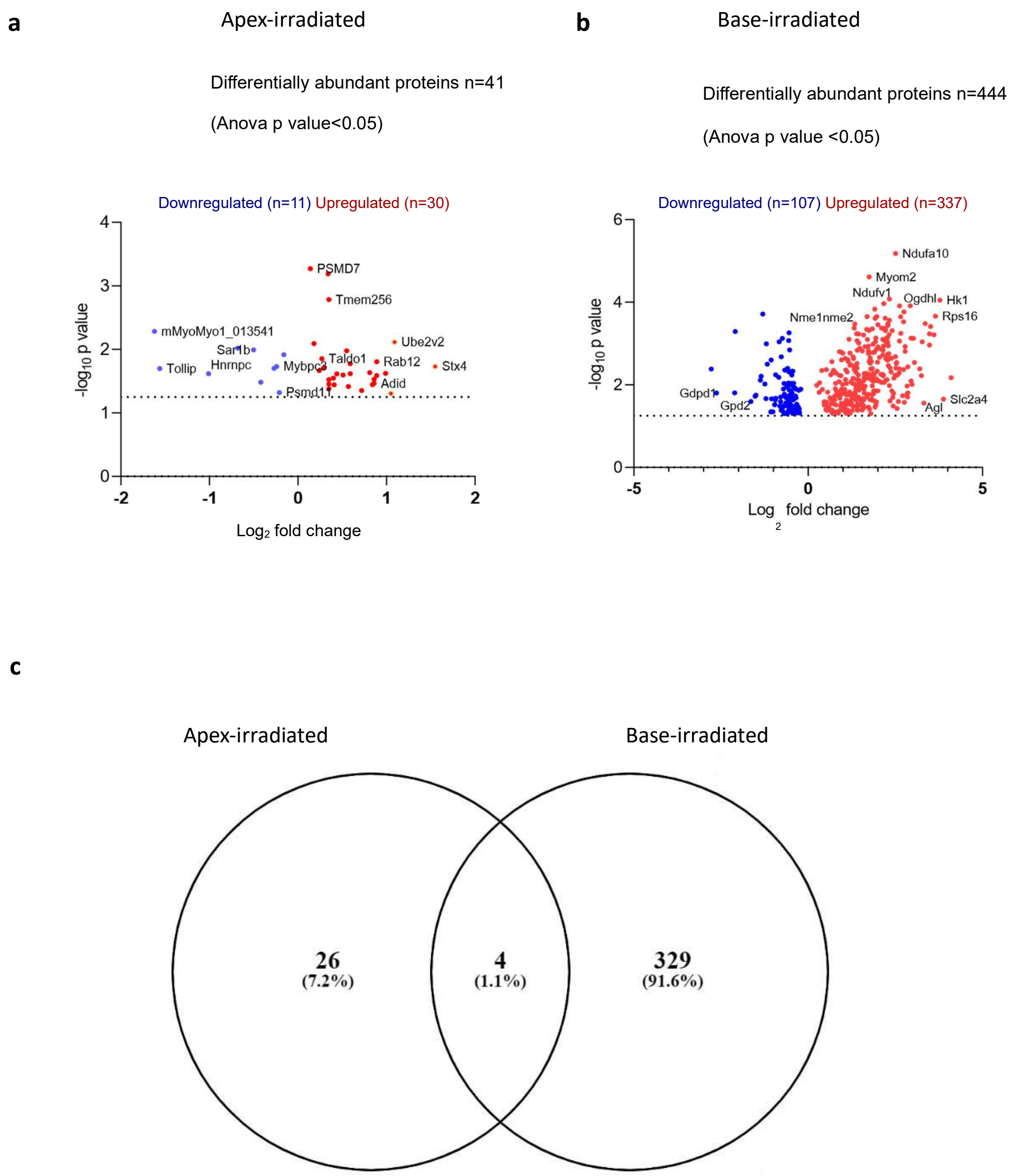


**Supplementary Figure 6. a,b** Proteomics-derived volcano plots of the downregulated (blue) and upregulated (red) proteins in either apex- (**a**) or base-irradiated (**b**) ventricle. –log10 (pvalue) cut-off was adopted to screen the statistically significant abundant proteins. **c**, Venny diagram illustrating the communal proteins observed during proteomics comparison of apex-irradiated and base-irradiated ventricular tissues. Venny 2.0.2 software was adopted to build the graph. Only proteins with associated gene nomenclature were adopted for this analysis.

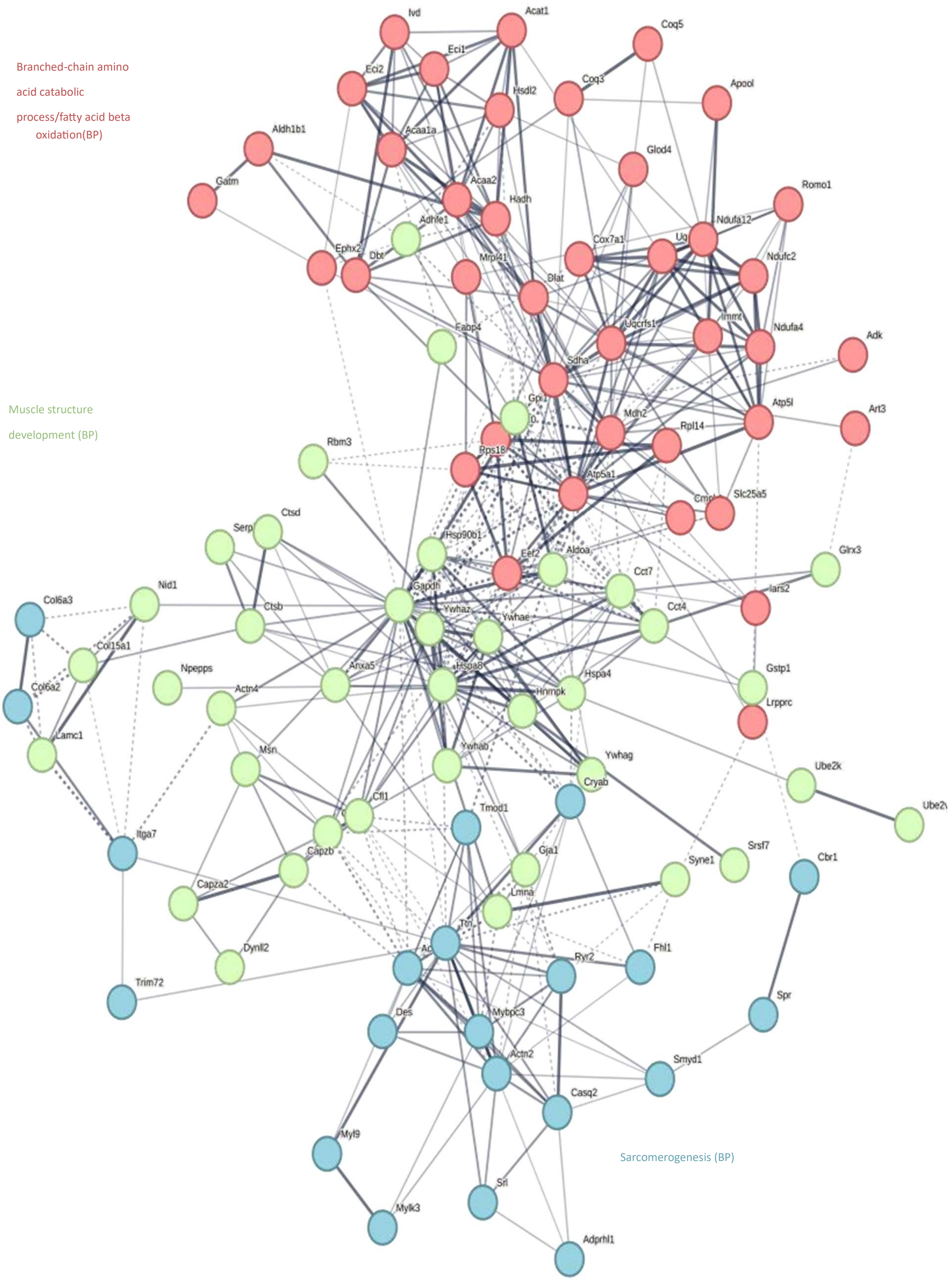

Branched-chain amino
acid catabolic
process/fatty acid beta
oxidation(BP)
Muscle structure
development (BP)
Sarcomerogenesis (BP)
Ivd
Acat1
Coq5
Eci1
Eci2
Coq3
Hsdl2
Apool
Aldh1b1
Acaa1a
Glod4
Gatm
Acaa2
Hadh
Romo1
Adhfe1
Ndufa12
Ephx2
Dbt
Mrpl41
Cox7a1
Ndufc2
Dlat
Fabp4
Uqcrfs1
Immt
Ndufa4
Adk
Sdha
Atp5l
Art3
Mdh2
Rpl14
Rbm3
Rps18
Atp5a1
Slc25a5
Ctsd
Serp
Hsp90b1
Eef2
Aldoa
Glrx3
Cct7
Gapdh
Iars2
Nid1
Col6a3
Ctsb
Ywhaz
Ywhae
Cct4
Col15a1
Npepps
Anxa5
Hspa8
Hspa4
Gstp1
Col6a2
Actn4
Hnrnpk
Lrpprc
Lamc1
Msn
Ywhab
Ywhag
Ube2k
Cryab
Cfl1
Tmod1
Itga7
Srsf7
Capzb
Gja1
Syne1
Cbr1
Capza2
Lmna
Ttn
Fhl1
Dynll2
Ryr2
Trim72
Des
Mybpc3
Spr
Actn2
Smyd1
Casq2
Myl9
Srl
Mylk3
Adprhl1

**Supplementary Figure 7.** K-Mean analysis of top 100 upregulated proteins in base-irradiated tissue by STRING 12.0 software form proteomics analysis 20weeks after radiation. Identification of functional patterns was performed with BP. KEGG or Reactome databases were adopted for the classification in case of missing BP identification, as reported in figure.

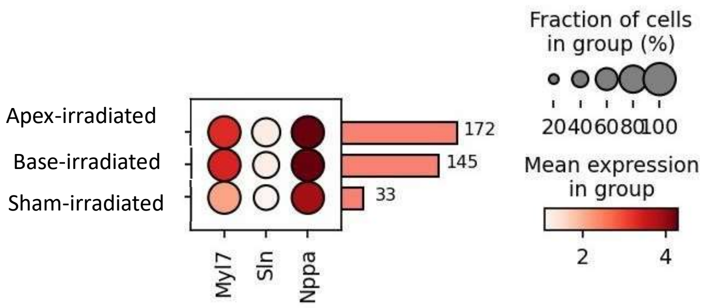


**Supplementary Figure 8. ST cardiomyocytes marker of right atrium.** Dot plot of ST of established atrial markers demonstrated preserved cardiomyocytes population despite radiation. Red bar on the right reported the number of analysed cells/sample in the ventricular clusters. Expression level is represented by the bar on the right.

a

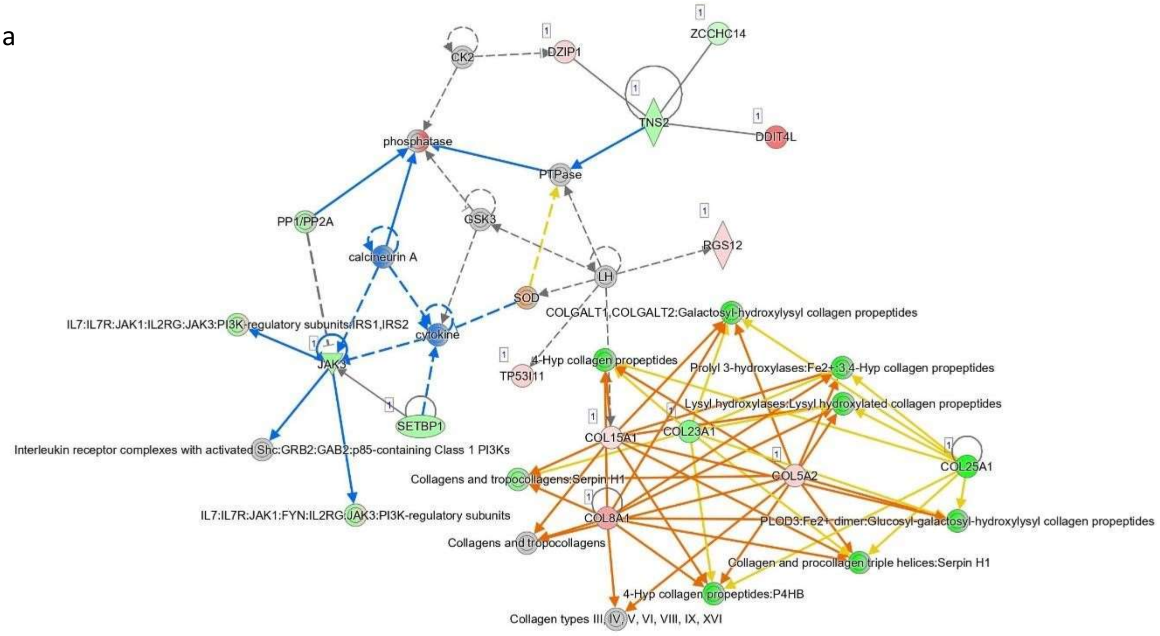


© 2000-2024 QIAGEN. All rights reserved.

b

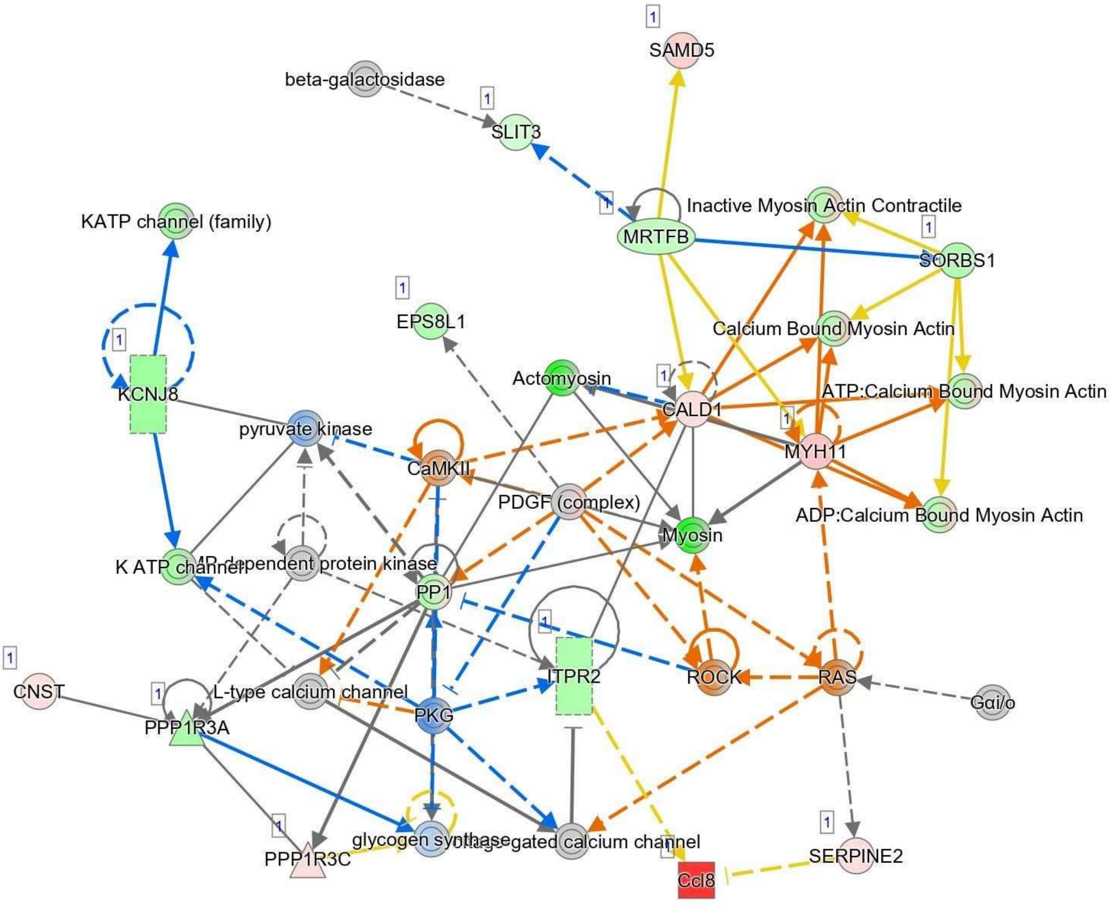


© 2000-2024 QIAGEN. All rights reserved.

c

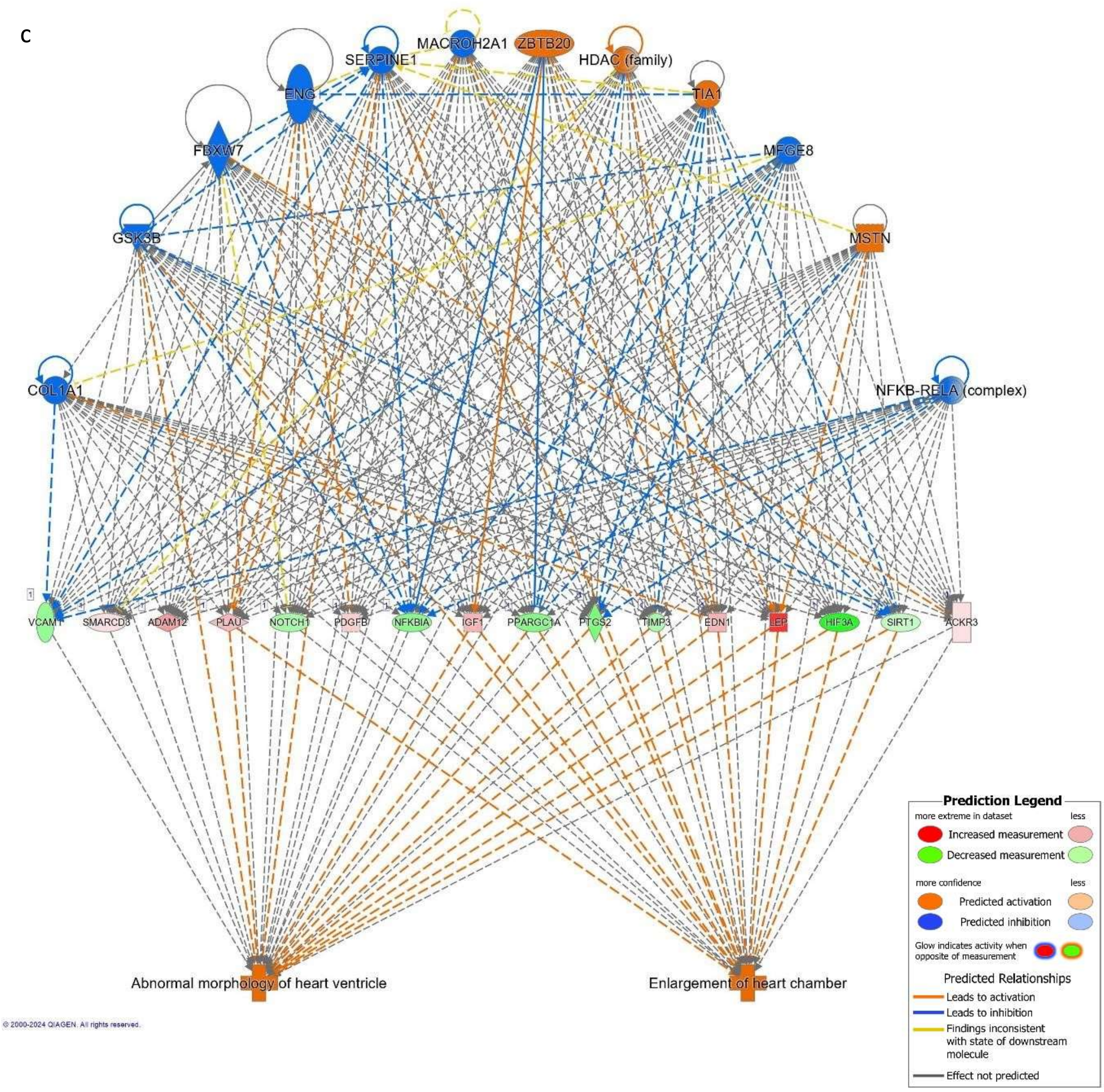


**Supplementary Figure 9. RNA-Sequencing analysis of right atrium using IPA.** Ingenuity Pathway Analysis (IPA) network of genes from RNA-Sequencing filtered using pAdjusted <0.1 dysregulated between sham-irradiated and right atrium tissue from base-irradiated tissue (**a,b**). Network, algorithmically generated based on their functional and biological connectivity, was graphically represented as nodes (genes) and edges (the biological relationship between genes). Expression of genes is reported according to the legend. Solid and dashed lines between genes represent known direct and indirect gene interactions, respectively. The shapes of the nodes reflect the functional class of each gene product: transcriptional regulator (horizontal ellipse), transmembrane receptor (vertical ellipse), enzyme (vertical rhombus), cytokine/growth factor (square), kinase (inverted triangle) and complex/group/other (circle). **a**: Collagen-related genes; **b**: conduction system-related genes. **c**, A network was generated to identify potential regulators of the major pathways within the right atrium using Upstream Regulator Analysis (URA). Interestingly, the final effect is highlighted as enlargement of the heart chamber and abnormal morphology.

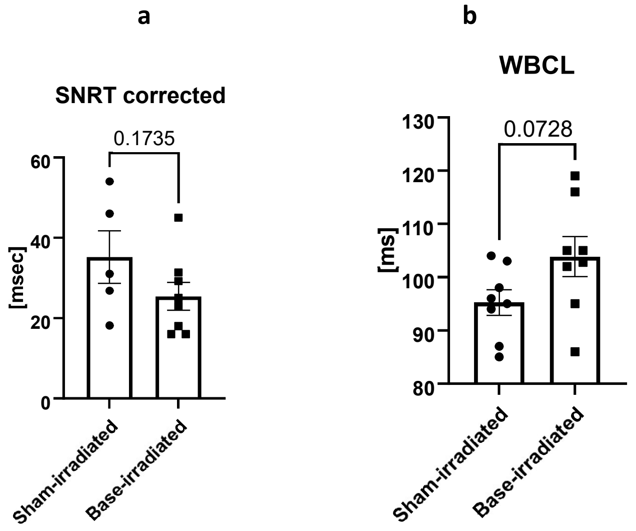


**Supplementary Figure 10. Ex vivo pacing parameters quantification. a,** Corrected SNRT quantification. Data are represented as mean±SEM and unpaired t-test was adopted. P value is reported in the graph. Sham-irradiated: n=6; base-irradiated: n=8. **b,** WBCL was quantified after appropriate pacing. WBCL data were normally distributed and unpaired t-test was adopted. Data are represented as mean±SEM. Sham-irradiated: n=8; base-irradiated: n=8. SNRT: Sinus Node Recovery Time. WBCL: Wenckebach cycle length.

| | Week 1 | | | Week 5 | | | Week 10 | | | Week 20 | | |
|---|---|---|---|---|---|---|---|---|---|---|---|---|
| | Sham-irradiated | Apex-irradiated | Base-irradiated | Sham-irradiated | Apex-irradiated | Base-irradiated | Sham-irradiated | Apex-irradiated | Base-irradiated | Sham-irradiated | Apex-irradiated | Base-irradiated |
| HR [sec] | 442±15.5 | 443±13 | 452±11.6 | 509±23.1 | 505±16 | 507±10 | 450±17.8 | 527±27.3 | 489±25.1 | 510±24 | 542±18 | 528±21 |
| PR interval [sec] | 40.2±1.4 | 39.2+0.6 | 38±0.8 | 39.4±0.6 | 39.2±0.7 | 39.1±1 | 40.1±2 | 39.6±0.7 | 39.6±1.2 | 39±1.4 | 42±.6 | 41.2±2 |
| P duration [sec] | 12.6±0.6 | 12.8±2.1 | 10.1±0.5 | 11±0.6 | 11.4±0.5 | 11±0.8 | 12.3±0.8 | 12.2±1.6 | 12.7±2 | 10±0.2 | 11.1±0.5 | 9.1±0.8 |
| QRS interval [sec] | 10.4±0.8 | 9.6±0.5 | 9.1±0.2 | 9.9±0.5 | 9.8±0.5 | 8.7±0.3 | 10.1±0.4 | 10±0.2 | 8.6±0.3* | 10.4±0.4 | 10.6±0.3 | 9.9±0.4 |
| QTc [sec] | 169.8±4.5 | 173.1±4.3 | 167.8±3 | 179±3.4 | 175±3.9 | 180.4±2 | 175±3.2 | 180±2.5 | 176±4 | 176.9±3.5 | 178.7±3 | 181.5±4 |
| JT interval [sec] | 27.9±4.6 | 28.3±3.2 | 32.9±9.3 | 36.5±2.4 | 33.3±4 | 38.1±1 | 30.9±3.5 | 29.5±3.6 | 35.1±2.1 | 31.1±3.2 | 33.2±5 | 33.5±2.5 |

**Supplementary Table 1. Longitudinal quantification of ECG parameters**. ECG parameters were measured over the different time points as presented in the table. QT interval was corrected with Framingham formula (QTc). Two-way ANOVA or Multiple Mann-Whitney test was adopted according to data distribution. No statistical significance was observed in parameters between groups and timepoints. *=QRS interval significant reduction compared to sham-irradiated, as showed in main body of article. HR= Heart rate.
